# Hypothalamic-extended amygdala circuit regulates temporal discounting

**DOI:** 10.1101/577973

**Authors:** Mark A. Rossi, Haofang E. Li, Glenn W. Watson, H. Gregory Moore, Min Tong Cai, Namsoo Kim, Katrina A. Vokt, Dongye Lu, Ryan A. Bartholomew, Henry H. Yin

**Author notes:** These authors contributed equally to this work (Mark A. Rossi, Haofang E. Li). Correspondence: Henry H. Yin.

## Abstract

Choice behavior is characterized by temporal discounting, i.e., preference for immediate rewards over delayed rewards. Temporal discounting is often dysfunctional in psychiatric disorders, addiction, and eating disorders. However, the underlying neural mechanisms governing temporal discounting are still poorly understood. We found that food deprivation resulted in steep temporal discounting of food rewards, whereas satiation abolished discounting. In addition, optogenetic activation of AgRP-expressing neurons in the arcuate nucleus or their axon terminals in the posterior bed nucleus of stria terminalis (BNST) restored temporal discounting in sated mice. Activation of postsynaptic neuropeptide Y receptors (Y1Rs) within the BNST, which is influenced by neuropeptide released by AgRP neurons, was sufficient to restore temporal discounting. These results demonstrate for the first time a profound effect of motivational signals from hypothalamic feeding circuits on temporal discounting and reveal a novel neural circuit that regulates choice behavior.

## Introduction

In choice behavior, the preference for an immediate over delayed reward is known as temporal or delay discounting. This preference for immediate gratification characterizes choice behavior in all species examined (Chung and Herrnstein, 1967; Green and Myerson, 2004; Kable and Glimcher, 2007). It reveals an evolutionarily conserved behavioral strategy that efficiently meets ongoing biological demands. In humans, however, it is often important to forego immediate rewards because they may have negative consequences. Aberrant temporal discounting is often associated with psychiatric disorders (Story et al., 2015), addiction (Bickel and Marsch, 2001), and obesity (Volkow et al., 2011). However, the underlying mechanisms of temporal discounting remain poorly understood. In particular, little is known about how discounting is modulated by intrinsic motivational states. The valuation of specific commodities, as reflected in choice behavior, should reflect the current motivational state and the status of essential homeostatic variables. For example, studies have shown that obesity is associated with greater temporal discounting, suggesting that higher demand for food is associated with a tendency for immediate gratification (Weller et al., 2008). However, the literature on choice behavior and decision making has not been integrated with the physiology of motivation.

Recent work in mice has begun to elucidate the neural mechanisms underlying intrinsic motivational states like hunger and thirst (Betley et al., 2013; Chen and Knight, 2016; Zimmerman et al., 2017). Can signals from these networks bias decision making and modulate temporal discounting? And if so, which signals are critical? One candidate originates from the group of neurons located within the arcuate nucleus of the hypothalamus (ARC) that express agouti-related peptide (AgRP) and neuropeptide Y (NPY). AgRP neurons monitor internal energy states by sensing circulating metabolic signals. They are activated by circulating hormones representing energy deficit, and inhibited by signals representing energy surfeit (Sternson, 2013). Moreover, AgRP neurons have been shown to be both necessary (Gropp et al., 2005; Luquet et al., 2005) and sufficient (Aponte et al., 2011; Krashes et al., 2011) for feeding in adult mice (Atasoy et al., 2012; Wu et al., 2009). Together, these observations suggest that AgRP neurons play a key role in signaling hunger within the brain, which is an integral component of the decision to eat. While previous studies have elucidated how AgRP neurons contribute to energy homeostasis, whether and how their outputs ultimately influence decision making remains unresolved (Rossi and Stuber, 2018).

We hypothesized that hunger-related signals from the AgRP neurons promote preference for immediate reward in choice behavior. Here, we showed for the first time that satiety abolished temporal discounting in mice, but selective activation of AgRP neurons can restore temporal discounting. AgRP neurons send projections to many subcortical brain regions including the bed nucleus of stria terminalis (BNST) (Betley et al., 2013), a key component of the extended amygdala that influences stress and anxiety (Chen et al., 2016; Pleil et al., 2015), food intake (Betley et al., 2013; Jennings et al., 2013), and choice behavior (Cullinan et al., 2008; Davis and Whalen, 2001) and is highly interconnected with brain regions associated with decision making and cognitive processing (Berthoud, 2007; O’Connell and Hofmann, 2012). We found that activation of AgRP axon terminals in the posterior BNST is sufficient to restore temporal discounting. This effect is due to the activation of postsynaptic neuropeptide Y1 receptors (Y1Rs) on target BNST neurons.

## Results

### Satiety abolishes temporal discounting in mice

Studies often measure temporal discounting as an approximation of self control or impulsivity using a design in which one choice leads to an immediate reward, while an alternate choice leads to a larger, delayed reward (Bickel and Marsch, 2001). This design allows researchers to test whether an animal is willing to forgo a delayed but larger reward for immediate gratification (Green and Myerson, 2004). However, using larger rewards for the delayed option can introduce a confound in studying pure delay discounting by manipulating both delay and reward size simultaneously. In the experiments described below, we utilize both approaches to probe the role of motivational state in guiding choice behavior and to elucidate its underlying mechanisms.

In order to assess pure temporal or delay discounting, we first designed an operant behavioral task, which uses the same reward size for both choices (Chung and Herrnstein, 1967). Mice were presented with two levers on each trial (**Figure 1A**). Pressing one lever always led to immediate food reward (14 mg pellet), whereas pressing the other lever led to delivery of the same food reward after a predetermined delay (0, 4, 8, or 16 s). We found that motivational state is a major determinant of temporal discounting. When mice were hungry, the discounting function was steep, and immediate rewards were robustly preferred over delayed rewards. However, when hungry mice were pre-fed immediately prior to testing or when mice were sated, they became indifferent to the delay (**Figure 1B**). Discounting functions were fit with a hyperbolic model in which the discounting parameter, *K*, indicates the steepness of the curve and thus the degree to which immediate rewards are preferred (Doya, 2008). Temporal discounting was greater when mice were hungry compared to when they were sated or pre-fed (**Figure 1C**, body weights shown in **Figure 1D**).

**Figure 1.**
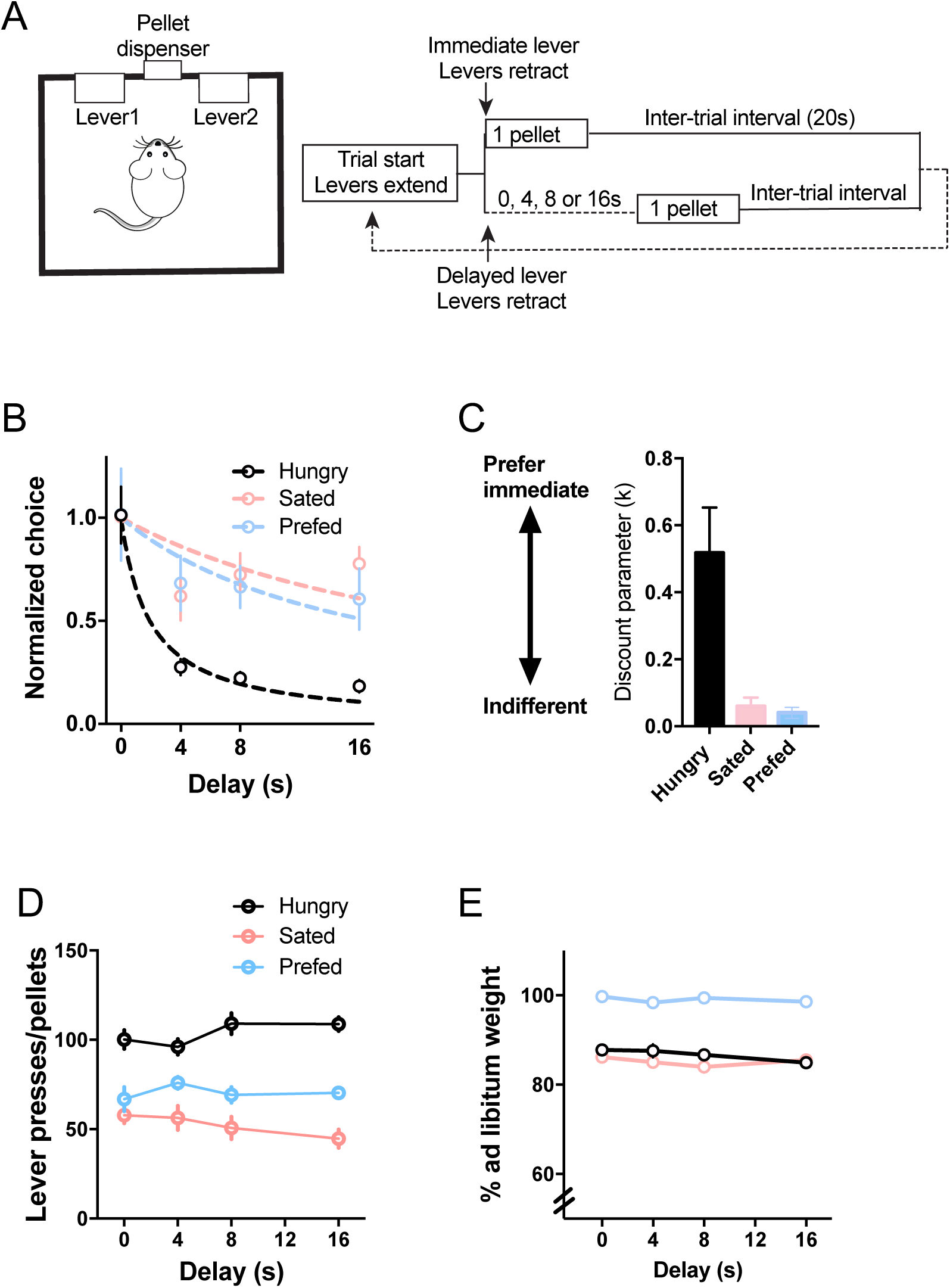
Temporal discounting depends on motivational state. **A.** Schematic of temporal discounting task. During each session, mice experienced only 1 delay. Mice were trained for 5-8 daily sessions on each delay. Temporal discounting was only measured after their performance had reached steady state. To control for biases in lever preference, the delay and non-delay levers were switched for each delay condition. **B.** Choice behavior varies as a function of motivational state. When mice are hungry (n = 13), immediate rewards are preferred, but when sated (n = 8) or following pre-feeding (n = 13), delayed rewards are discounted less. Dashed lines are hyperbolic fits using the equation: Y = 1/(1+*K*\*d). Choice of the delay lever (Y) varies as a function of delay, d, and the discounting parameter, *K*, which determines the steepness of discounting. To obtain the normalized choice measure, we divided number of presses on the delay lever by the baseline number when there is no delay. **C.** Discounting parameters corresponding to hyperbolic fits in B. The higher the value of *K*, the greater the preference for the immediate reward (steeper discounting function). A one-way ANOVA showed a main effect of satiety (F _2, 31_ = 9.36, p = 0.0007, Tukey’s multiple comparisons test: Sated vs. Hungry, p = 0.0012; Sated vs. Prefed, p = 0.99; Hungry vs. Prefed, p = 0.0066). Values are mean and s.e.m. **D.** Total number of presses/rewards per session. A two-way ANOVA showed no interaction between motivational state and delay (F _6, 121_ = 1.63, p = 0.14,) no significant main effect of delay (F _3, 121_ = 0.069, p = 0.98), and a significant main effect of motivational state (F_2, 121_ = 104.1, p < 0.0001). **E.** Body weights for all mice tested.

### AgRP activation restores temporal discounting in sated mice

AgRP neurons are necessary for voluntary feeding (Gropp et al., 2005; Luquet et al., 2005), respond to changing metabolic needs (Sternson, 2013), and can influence many motivated behaviors (Burnett et al., 2016; Dietrich et al., 2015; Padilla et al., 2016), but their role in decision making is unknown. Thus, we aimed to test whether AgRP activation biases temporal discounting. To selectively activate AgRP neurons *in vivo*, Ai32 mice, which express a floxed-STOP codon and ChR2-eYFP under the ROSA26 locus (Madisen et al., 2012), were crossed with AgRP-Cre mice (Tong et al., 2008), yielding ChR2 expression restricted to AgRP neurons (AgRP::ChR2; **Figures 2A-C**, and **Extended Figure 1**). To validate the efficacy of ChR2 within AgRP neurons, whole-cell patch clamp electrophysiology was performed in acute brain slices from adult AgRP::ChR2 mice. Blue light pulses result in inward currents in voltage clamp recordings and reliable spiking in current clamp recordings (**Figure 2D**).

**Figure 2.**
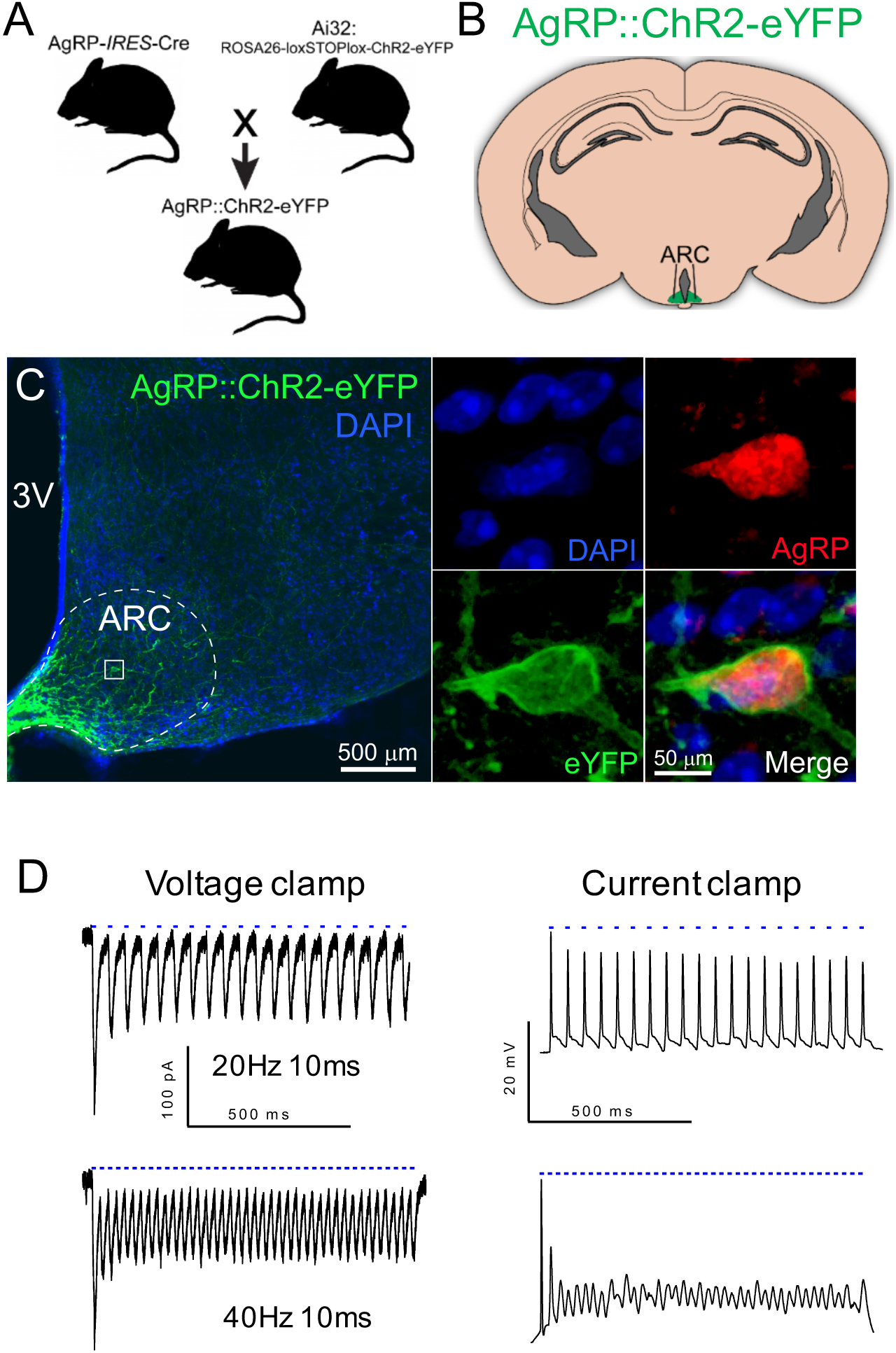
Genetic targeting of AgRP neurons with ChR2. **A.** Scheme for generating AgRP::ChR2 mice. **B.** Schematic representation of restricted ChR2 expression within the ARC. **C.** Representative expression pattern of AgRP-ChR2-eYFP within the ARC. Scale bar, 50 µm. Transgenic expression of ChR2 results in membrane-bound eYFP and nuclear exclusion. 3V = third ventricle. ARC =arcuate nucleus. **D.** Whole-cell patch clamp recordings from ChR2-expressing ARC neurons. 10 ms blue light pulses result in inward currents in voltage clamp recordings and spiking in current clamp recordings. Top row, 20 Hz stimulation; bottom row, 40 Hz stimulation.

We then tested the hypothesis that optogenetic activation of AgRP neurons is sufficient to restore temporal discounting in sated mice. We targeted the ARC of AgRP::ChR2 mice with optic fibers (AgRP::ChR2^ARC^; **Figure 3A**) to allow delivery of 1 s pulsed trains of 473 nm laser light (1 s pulsed at 10, 20, or 40 Hz followed by three seconds of no stimulation repeated for the duration of the experiment). Mice with chronically implanted optic fibers were trained on the temporal discounting task. Replicating our finding that motivational state biases decision making, hungry mice strongly preferred immediate food rewards to delayed rewards, but such temporal discounting was abolished by satiety (**Figure 3B** and **Extended Figure 2**). In sated mice, however, AgRP::ChR2^ARC^ stimulation restored discounting observed in the hungry state (see **Extended Figure 3** for representative examples).

**Figure 3.**
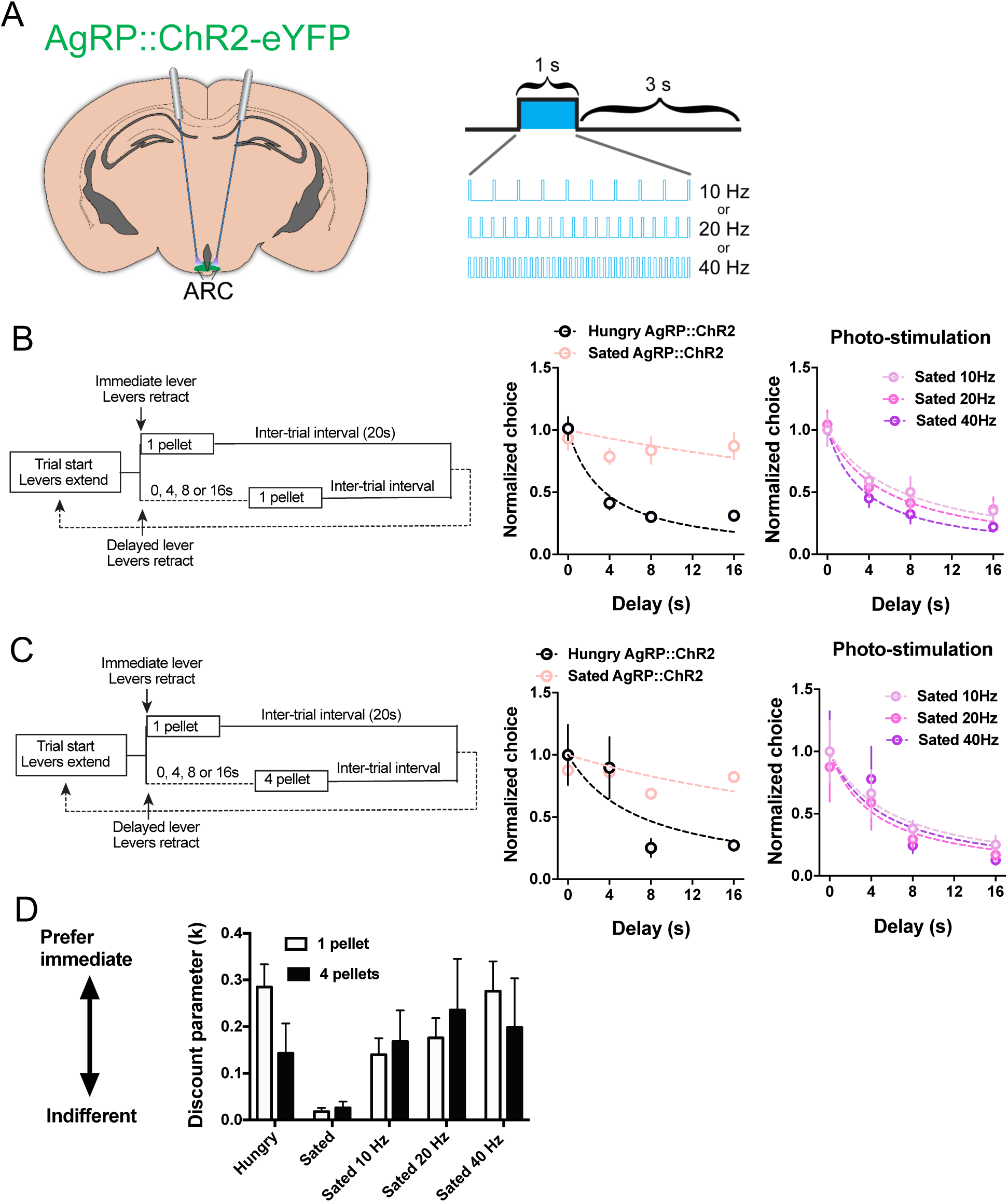
AgRP stimulation restores delay discounting. **A.** Bilateral optical fibers were implanted in AgRP::ChR2-eYFP mice targeting ARC. 1-s pulse trains were delivered every 4 seconds at 10 Hz, 20 Hz, or 40 Hz. **B.** Temporal discounting functions for AgRP::ChR2 mice in the pure delay discounting condition, in which the delayed reward is the same size as the immediate reward (1 pellet). Dashed lines represent hyperbolic fits. Discounting can be restored to Hungry levels by AgRP stimulation. Error bars indicate ± s.e.m. Hungry (n = 23), sated (n = 23), Sated 10 Hz (n =11), Sated 20 Hz (n = 23), Sated 40 Hz (n = 22). Values are mean and s.e.m. **C.** Discounting functions for the delay/larger reward (4 pellets) condition. **D.** Summary of the discounting parameter (*K*), which is a measure of temporal discounting, for different groups. AgRP::ChR2 mice show typical state-dependent discounting of delayed rewards. Satiety largely abolishes temporal discounting. However, AgRP::ChR2 stimulation restores discounting in sated mice. The reward size associated with the delay choice was either the same as the non-delay choice (1 pellet, pure delay discounting) or larger than the non-delay choice (4 pellets, delay/reward size tradeoff). A two-way ANOVA with stimulation frequency and reward size revealed no interaction (F _3, 104_ = 0.44, p = 0.73), a main effect of stimulation, F _3, 104_ = 4.51, p = 0.005), and no main effect of reward size (F _1, 104_ = 0.01, p = 0.92). Hungry (n = 8), Sated (n = 8), Sated 10 Hz (n =8), Sated 20 Hz (n = 8), Sated 40 Hz (n = 8).

In addition, we also tested the same mice using the delayed/larger reward design often used as a test of impulsivity and self control (Cardinal et al., 2001). Instead of delivering one pellet following the choice of the delayed lever, four pellets were delivered. A similar hyperbolic discounting function was observed (**Figure 3C**), but the larger reward slightly reduced the steepness of discounting when mice were hungry, confirming that regardless of the size of the delayed reward, AgRP::ChR2^ARC^ stimulation can restore discounting. The steepness of discounting, as quantified by the discount parameter (k), depended on the stimulation frequency (**Figure 3D**).

In addition to choice behavior, AgRP::ChR2^ARC^ stimulation increased the total number of lever presses (choices) and pellets earned per session, suggesting higher overall motivational drive acting independently of the choices made (**Figure 4B** and **Figure 4D**). This finding is consistent with previous studies demonstrating that AgRP activation increases willingness to work for food (Atasoy et al., 2012; Krashes et al., 2011). Laser stimulation failed to affect the choice behavior of AgRP-Cre littermates lacking functional ChR2 (ChR2-Cre control mice, **Extended Figure 4**). Nor did stimulation affect overall lever pressing in these mice (**Figure 4A** and **Figure 4C**). Taken together, these results indicate that, in addition to influencing food intake and competing innate behaviors, AgRP activation potently biases choice behavior toward impulsive decisions.

**Figure 4.**
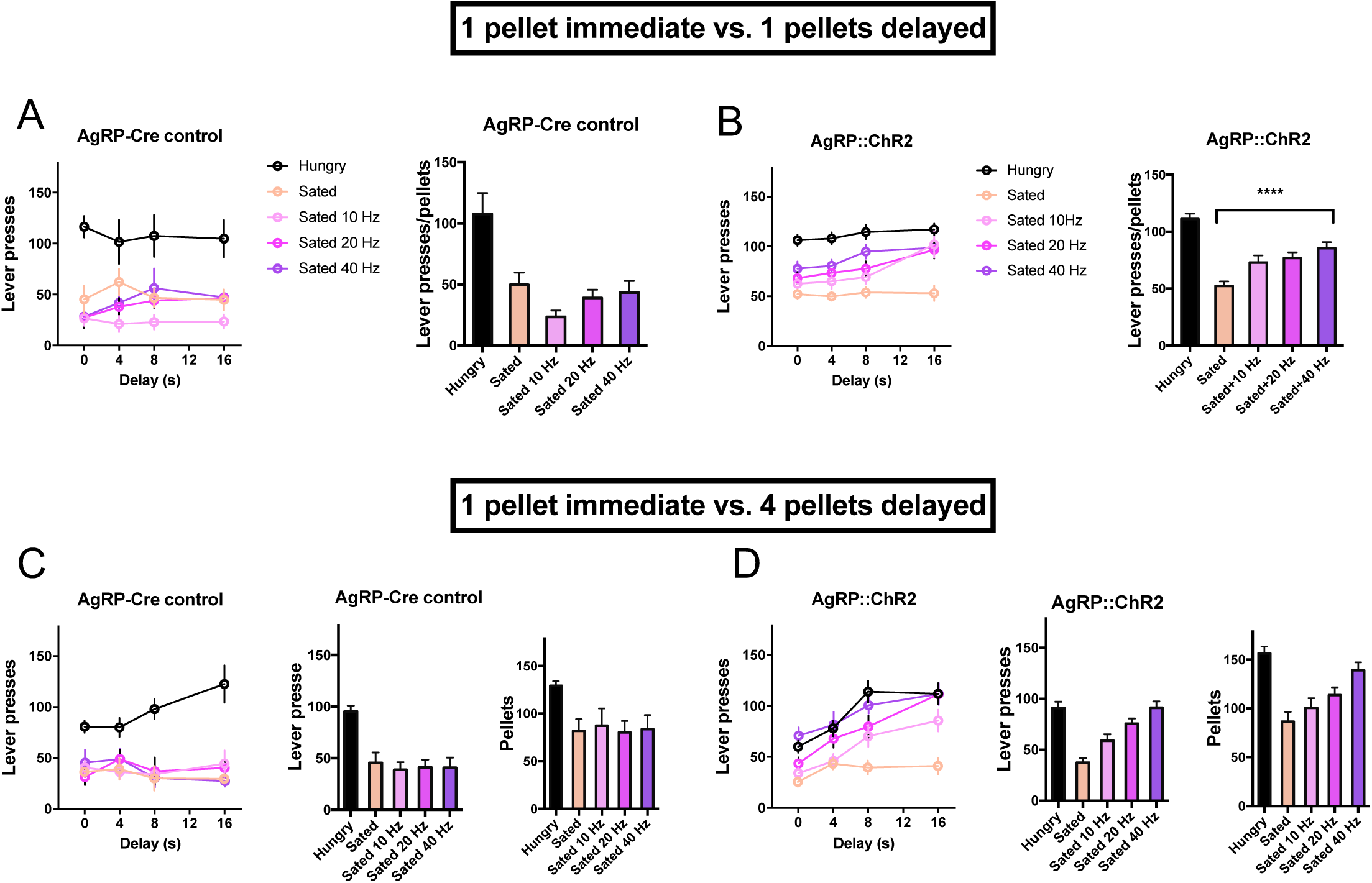
AgRP stimulation increases total lever pressing. **A.** Total number of lever presses (choices) per 1-hour session from AgRP-Cre control mice. Left, averaged lever pressing data from the last two sessions of each delay condition. Right, summary with total lever pressing averaged across different delays. In the AgRP-Cre controls, a one-way ANOVA on the optogenetic stimulation data (sated only) showed no main effect of stimulation (F _3, 19_ = 1.67, p = 0.21). The Hungry group showed higher lever pressing all other groups (Dunnett’s multiple comparisons test, p = 0.003, p = 0.0001, p = 0.0005, p = 0.001). Since each choice always yields one pellet, the number of total presses and number of pellets earned are identical. **B.** Total number of lever presses (choices) per 1-hour session from AgRP::ChR2 mice. There is a significant main effect of stimulation (F _3, 28_ = 9.23, p = 0.0002). Lever pressing was higher in the Hungry group compared to sated (p = 0.0001), and sated 10 Hz (p = 0.001), but at higher frequencies stimulation restored overall lever pressing to that of the Hungry group: 20 Hz (p = 0.07) or 40 Hz (p = 0.42). Hungry (n = 23), sated (n = 23), Sated 10 Hz (n =11), Sated 20 Hz (n = 23), Sated 40 Hz (n = 22). **C.** Similar results on total lever pressing were found in the larger delayed reward (4 pellets) condition. There was no effect of stimulation in AgRP-Cre control mice (F _3, 16_ = 0.10, p = 0.96). The Hungry group showed higher lever pressing all other groups (Dunnett’s multiple comparisons test, p < 0.0001 regardless of stimulation frequency). Because the delayed reward is larger, the number of pellets is plotted separately (right). **D.** AgRP::ChR2 mice also showed a significant effect of stimulation in the larger delayed reward condition (F _3, 18_ = 17.18, p < 0.0001). Comparing the Hungry condition with all others, there was a significant difference between Hungry and Sated (p = 0.0001), and Sated 10 Hz (p = 0.02), but at higher frequencies stimulation restored overall lever pressing to a level comparable to the Hungry group: 20 Hz (p = 0.56) or 40 Hz (p = 0.88). Because the delayed reward is larger, the total number of pellets earned per session is plotted separately (right). Hungry (n = 8), Sated (n = 8), Sated 10 Hz (n =8), Sated 20 Hz (n = 8), Sated 40 Hz (n = 8).

### AgRP projections to posterior BNST mediate temporal discounting

The BNST is a target of AgRP projections (Betley et al., 2013) that regulates stress and anxiety (Chen et al., 2016; Pleil et al., 2015), feeding (Betley et al., 2013; Jennings et al., 2013), and high-level choice behavior (Cullinan et al., 2008; Davis and Whalen, 2001). In addition, the BNST is connected with brain regions associated with decision making and cognitive processing, including the ventral tegmental area, hippocampus, and prefrontal cortex (Berthoud, 2007; O’Connell and Hofmann, 2012; Stamatakis et al., 2014). Thus, we aimed to test whether AgRP activation can influence temporal discounting. We stimulated the axon terminals of AgRP neurons in the BNST (**Figure 5A**). AgRP::ChR2→BNST stimulation restored discounting and increased lever pressing in sated mice (**Figure 5** and **Extended Figure 5**, see body weights and representative examples in **Extended Figures 6** and **7**). Photo-stimulation frequency-dependently restored discounting in both identitcal delayed reward (**Figure 5B**) and larger delayed reward (**Figure 5C**) conditions. Overall, the larger delayed reward produced less discounting (**Figure 5D**).

**Figure 5.**
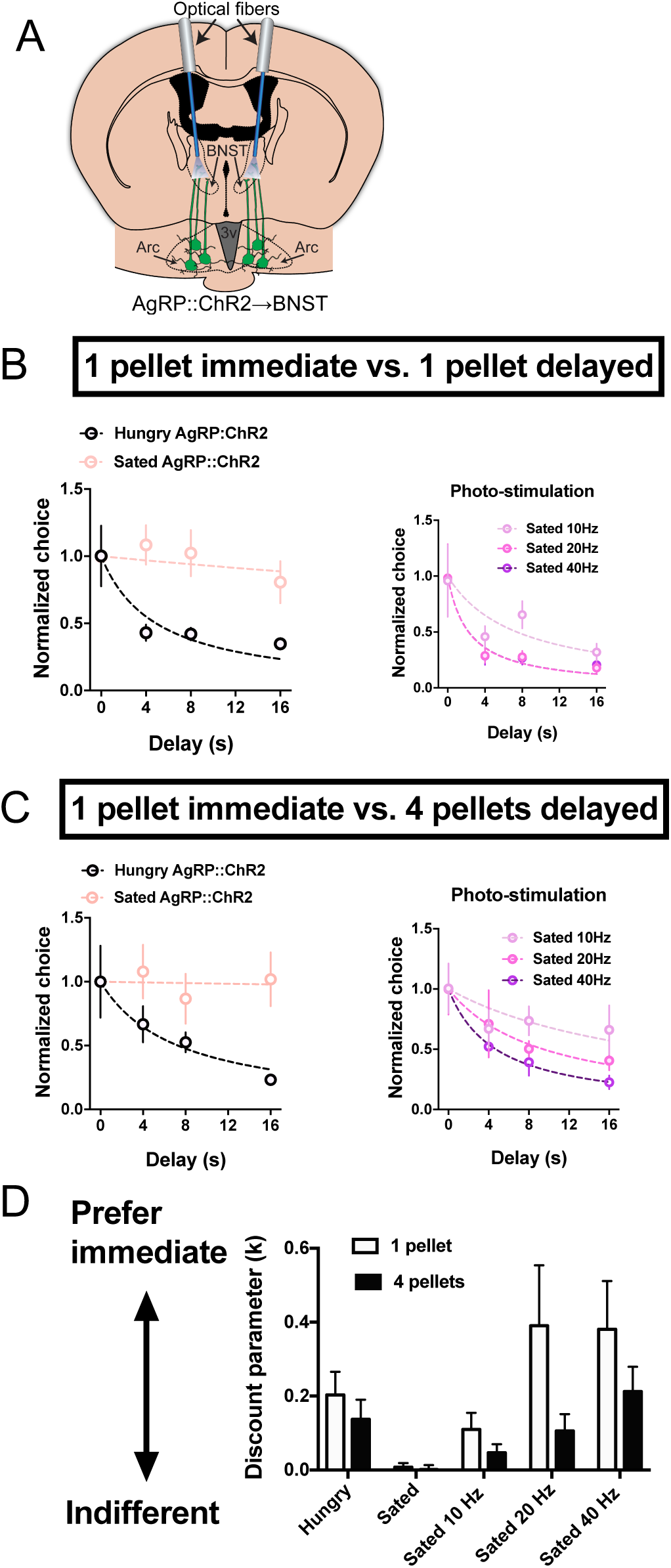
Stimulation of AgRP::ChR2 terminals in BNST restores temporal discounting in sated mice. **A.** Schematic illustration of AgRP terminal stimulation in BNST. Bilateral optical fibers were implanted in the posterior BNST of AgRP::ChR2-eYFP mice. **B.** Temporal discounting functions for AgRP::ChR2→BNST stimulation in the pure delay discounting condition, in which the delayed reward is the same size as the immediate reward (1 pellet). Dashed lines represent hyperbolic fits. Discounting can be restored to Hungry levels by AgRP stimulation. Hungry (n = 8), sated (n = 8), Sated 10 Hz (n =8), Sated 20 Hz (n = 8), Sated 40 Hz (n = 8). **C.** Discounting functions for the delay/larger reward (4 pellets) condition. **D.** Summary of discounting parameter (*K*), which is a measure of temporal discounting, for different groups. AgRP::ChR2 mice show typical state-dependent discounting of delayed rewards. Satiety largely abolished temporal discounting. However, AgRP::ChR2→BNST stimulation restores discounting in sated mice. The reward size associated with the delay choice was either the same as the non-delay choice (1 pellet, pure delay discounting) or larger than the non-delay choice (4 pellets, delay/reward size tradeoff). A two-way ANOVA with stimulation frequency and reward size revealed no interaction (F _3, 52_ = 1.01s, p = 0.39), a main effect of stimulation, F _3, 52_ = 5.14, p = 0.0034), and a main effect of reward size (F _1, 52_ = 4.61, p = 0.036). Thus, while stimulation frequency-dependently restored discounting in both the same delayed reward and larger delayed reward conditions, overall the larger delayed rewards produced less discounting. Hungry (n = 7), Sated (n = 7), Sated 10 Hz (n =7), Sated 20 Hz (n = 7), Sated 40 Hz (n = 7).

**Figure 6.**
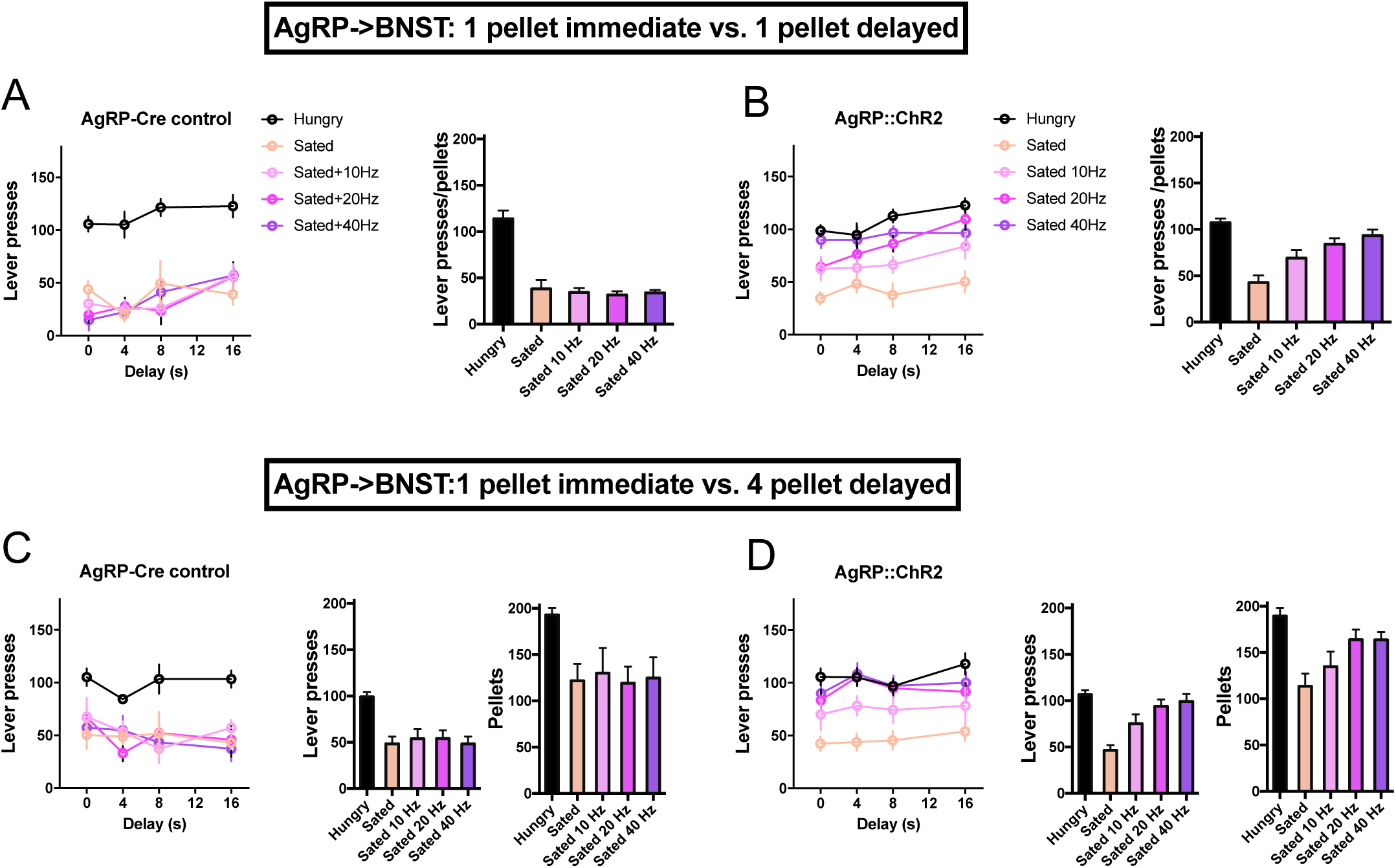
Summary of lever pressing in AgRP→BNST terminal stimulation. **A.** Total number of lever presses (choices) per 1-hour session from the AgRP-Cre control mice. Left, averaged lever pressing data from the last two sessions of each delay condition, once performance has reached steady state. Right, summary with total lever pressing averaged across different delays. In the AgRP-Cre controls, a one-way ANOVA on the optogenetic stimulation data (sated only) showed no main effect of stimulation (F_3, 16_ = 0.20, p = 0.89). The Hungry group showed higher lever pressing than all other groups (Dunnett’s multiple comparisons test, p < 0.0001) regardless of stimulation frequency. **B.** Total number of lever presses (choices) per 1-hour session from AgRP::ChR2 mice. There is a significant main effect of stimulation (F _3, 28_ = 9.23, p = 0.0002). Lever pressing was higher in the Hungry group compared to Sated (p = 0.0001), and Sated 10 Hz (p = 0.001), but at higher frequencies stimulation restored overall lever pressing: 20 Hz (p = 0.07) or 40 Hz (p = 0.42). Hungry (n = 8), Sated (n = 8), Sated 10 Hz (n =8), Sated 20 Hz (n = 8), Sated 40 Hz (n = 8). **C.** Similar results on total lever pressing were found in the larger delayed reward (4 pellets) condition. There was no effect of stimulation in AgRP-Cre control mice (F _3, 16_ = 0.13, p = 0.94). The Hungry group showed higher lever pressing compared to all other groups (Dunnett’s multiple comparisons test, p < 0.0001 for all stimulation frequencies). Because the delayed reward is larger, the total number of pellets earned per session is plotted separately (right). **D.** Total number of lever presses (choices) per 1-hour session from AgRP::ChR2 mice (F _3, 24_ = 9.22, p < 0.0003, p = 0.0001). Comparing the Hungry group with all others, there was a significant difference in lever pressing between Hungry and Sated (p = 0.0001), and Sated 10 Hz (p = 0.02). However, at higher frequencies stimulation restored lever pressing to a level comparable to that of the Hungry group: (20 Hz, p = 0.56, 40 Hz, p = 0.88). Because the delayed reward is larger, the total number of pellets earned per session is plotted separately (right). Hungry (n = 7), Sated (n = 7), Sated 10 Hz (n =7), Sated 20 Hz (n = 7), Sated 40 Hz (n = 7).

**Figure 7.**
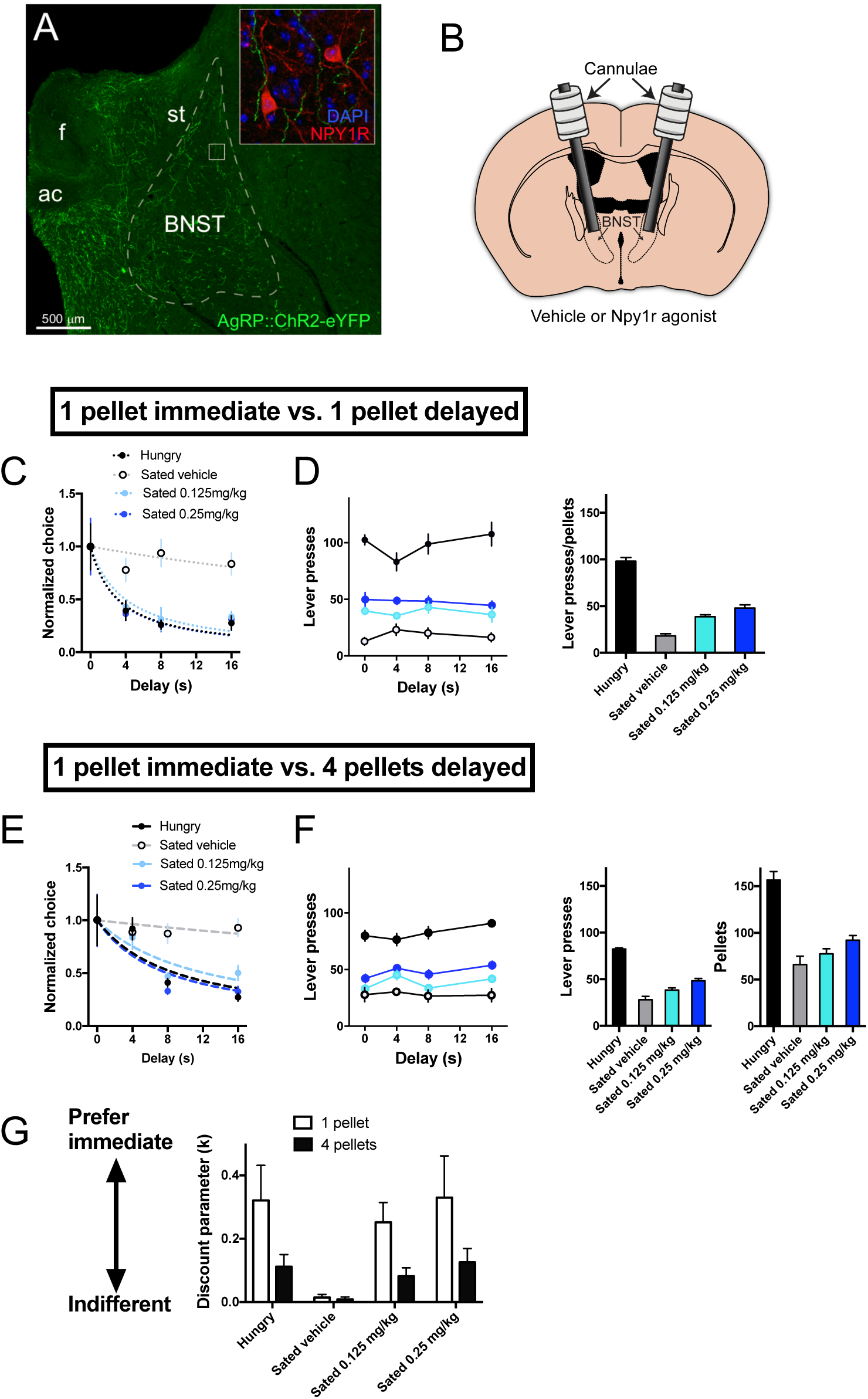
BNST^Y1R^ activation is sufficient for temporal discounting. **A.** Axons of AgRP::ChR2-eYFP neurons target postsynaptic BNST neurons that express neuropeptide Y 1 receptors (NPY1Rs). Ac = Anterior commissure, BNST = bed nucleus of the stria terminalis, F = fornix, St = stria terminalis. **B.** Schematic illustration of bilateral cannulae chronically implanted into the BNST. **C.** Temporal discounting functions for Y1R agonist experiments using the same reward size for the delayed lever (1 pellet). Dashed lines represent hyperbolic fits. **D.** Total number of lever presses (choices) per 1-hour session for the same size delayed reward condition. Left, averaged lever pressing data from the last two sessions of each delay condition. Right, summary with total lever pressing averaged across different delays. A one-way ANOVA on the drug injection results (sated only) showed a main effect of dose (F _2, 15_ = 32.9, p < 0.0001). The Hungry group showed higher lever pressing than all other groups (Dunnett’s multiple comparisons test, ps = 0.0001). Thus, while Y1R activation significantly increased lever pressing, it did not restore it to the level seen in the Hungry group. **E.** Temporal discounting functions using a larger reward size for the delayed lever (4 pellets). Dashed lines represent hyperbolic fits. Hungry (n = 6), Sated Vehicle (n = 6), Sated 0.125 mg/kg (n = 6), Sated 0.25 mg/kg (n =6). **F.** Total number of lever presses (choices) per 1-hour session for the larger delayed reward condition. Left, averaged lever pressing data from the last two sessions of each delay condition. Right, summary with total lever pressing averaged across different delays. A one-way ANOVA on the drug injection results (sated only) showed a main effect of dose (F _2, 15_ = 12.30, p = 0.0007). The Hungry group showed higher lever pressing than all other groups (Dunnett’s multiple comparisons test, p = 0.0001 for all comparisons). Thus, while Y1R activation significantly increased lever pressing, it did not restore it to the level seen in the Hungry group. Because the delayed reward is larger, the total number of pellets earned per session is plotted separately (right). **G.** Discounting parameter summary for Y1R agonist (Leu-Pro NPY) experiments. Activation of Y1R generated delay discounting in sated mice. A two-way ANOVA showed no significant interaction between dose and reward size (F _3, 40_ = 0.97, p = 0.41), a significant main effect of drug (F _3, 40_ = 4.20, p = 0.013), and a significant main effect of reward size (F _1, 40_ = 9.21, p = 0.004). Thus, Y1R activation dose-dependently restored discounting. In addition, the degree of discounting is reduced in the larger delayed reward condition. Error bars indicate ± s.e.m.

In addition, AgRP::ChR2→BNST stimulation also restored the number of lever presses and rewards earned (**Figure 6B,D**). At higher stimulation frequencies, the rate of lever pressing and pellets earned were promoted to a level comparable to that of the Hungry group. In AgRP-Cre control mice, discounting and lever pressing are unaffected by stimulation (**Figure 6A,C** and **Extended Figure 8**).

### NPY signaling in the posterior BNST is critical for temporal discounting

We next sought to probe the underlying pharmacological mechanism driving the observed changes in temporal discounting. AgRP neurons co-express the orexigenic neuropeptide, NPY (Hahn et al., 1998), and NPY receptor activation contributes to some AgRP-mediated behaviors (Atasoy et al., 2012; Dietrich et al., 2015). As shown in **Figure 7A**, AgRP axons in the BNST target neurons that express type 1 NPY receptors (Y1Rs), providing a potential mechanism for motivational signals to impact downstream structures mediating choice behavior. To test whether BNST^Y1R^ activation is sufficient to control temporal discounting, we employed a pharmacological approach. Mimicking the effects of Y1R activation, we injected the high affinity Y1R agonist Leu-Pro NPY into the BNST region that receives AgRP projections (**Figure 7B** and **Extended Figure 9**). BNST^Y1R^ activation is sufficient to restore temporal discounting in sated mice (**Figure 7C** and **Figure 7E**; see **Extended Figure 10** for body weights and representative examples). BNST^Y1R^ activation also significantly increased total lever pressing per session, but it did not restore it to the level seen in the Hungry group (**Figure 7D** and **Figure 7F**). The larger delayed reward condition also produced less discounting overall, as expected, but BNST^Y1R^ activation significantly increased temporal discounting regardless of the size of the delayed reward (**Figure 7G**).

## Discussion

Our results revealed a novel neural pathway contributing to reward-guided decision making. We found that the AgRP→BNST projection strongly modulates choice behavior. Temporal discounting is largely abolished in sated mice, but is restored by activation of AgRP neurons in the ARC or their axon terminals in the posterior BNST, a major downstream target. As AgRP neurons are known to express NPY, we tested whether activation of postsynaptic NPY receptors (Y1Rs) on BNST neurons is sufficient to restore temporal discounting. Similar to optogenetic stimulation of AgRP cell bodies or axon terminals within the BNST, BNST^Y1R^ activation biases behavior toward choosing immediate rewards. Although BNST^Y1R^ activation dose-dependently increased temporal discounting, it did not restore discounting to the levels of hungry mice, which was the case for optogenetic activation. This suggests that in addition to NPY, AgRP neurons may regulate choice behavior through co-release of other transmitters, such as AgRP or GABA (Atasoy et al., 2012; Tong et al., 2008; Wu et al., 2009).

### Motivational modulation of temporal discounting

Our behavioral results demonstrate that motivational state potently modulates temporal discounting. Hungry mice strongly preferred immediate food rewards, yet sated mice were largely indifferent to reward delay. This is true whether the delayed reward is the same size (pure delay discounting) or much larger than the immediate reward (self control). Delay discounting is by definition discounting of the delayed reward (Chung and Herrnstein, 1967). The delay/reward size tradeoff design was introduced primarily to study the question of ‘self control,’ which is concerned with whether one should choose a small reward now versus big reward later (Green and Myerson, 2004). We observed the same pattern of behavior when a larger reward was associated with the delayed choice. Increasing reward size for the delayed lever, as expected, merely reduced the steepness of discounting. Moreover, AgRP cell body stimulation or AgRP→BNST terminal stimulation both restored discounting whether or not the delayed reward is larger.

An obvious question is why mice should choose the delayed option at all when the delayed reward is the same size as the immediate reward. It should be noted that, because environmental contingencies are not always fixed in nature, some sampling of the currently less desirable option is an explorative strategy that allows the animal to update their representation of action-outcome contingencies. Our results show that this is modulated by motivational state, in agreement with previous work showing that satiety also results in indifference to risk (variance in desired rewards): when mice are faced with the choice of certain reward and uncertain reward, they prefer certain rewards when hungry, yet become indifferent when sated (Leblond et al., 2011; Leblond et al., 2014).

Previous studies in pigeons and humans have examined how motivational state influences choice behavior, but no clear consensus was established, in part due to the experimental design used (Kirk and Logue, 1997; Logue and Pena-Correal, 1985). They always used the self control design in which larger rewards are associated with the delayed lever (Rachlin and Green, 1972), which may introduce confounds. Moreover, previous work that failed to see effects of motivational state on choice behavior used the same lever for the delayed choice (Logue and Pena-Correal, 1985). This design could promote a strong bias for exclusive choice of the immediate lever. Due to the lack of sampling of the other lever, animals never learn about the actual delays associated with each lever. On the other hand, in our experiments, we specifically controlled for the lever or location bias. We switched the lever-delay assignment after changing to a different delay and trained the animals until their choice performance reached steady state. Thus, our design ensures that delay is the only factor determining choice behavior, not lever location or any other previously learned bias.

It should also be noted that, in all experiments, mice ate most or all of the pellets that they earned. Despite satiation, mice continued to earn and consume food rewards, though at a lower rate than when mice are hungry. The most likely explanations for why mice would continue to work for and consume food when sated is that 1) the mice are experienced with food restriction, which is known to promote overeating (Hagan et al., 2002), and 2) motivation to eat is potentiated by the palatability of the food pellets relative to their home chow. Palatability is well known to potentiate food intake (Berridge, 1996). Future studies will be needed to understand the specific contributions of the history of food restriction and palatability to AgRP-mediated temporal discounting.

### The role of AgRP signaling in choice behavior

By optogenetically activating AgRP neurons in the ARC, we were able to restore temporal discounting in sated mice (**Figure 3**). Stimulation frequency determines *K*, the parameter representing the steepness of discounting. This is to our knowledge the first report of a causal role of a genetically defined neuronal population in temporal discounting (for a comprehensive review, see (Frost and McNaughton, 2017). Interestingly, AgRP cell body stimulation restored temporal discounting in sated mice regardless of the reward size associated with the delayed lever. The fact that AgRP activation can supersede self control is in agreement with recent studies demonstrating that AgRP activation competes with and can override other motivated behaviors associated with aggression, anxiety, fear, social interaction, grooming, locomotion, foraging, and reward (Burnett et al., 2016; Dietrich et al., 2015; Padilla et al., 2016).

Recent work showed that AgRP neurons are rapidly inhibited by food or cues that predict food (Betley et al., 2015; Burnett et al., 2016; Chen et al., 2015; Mandelblat-Cerf et al., 2015). On the other hand, AgRP activation appears to initiate food consumption, including instrumental behaviors directed at food (Atasoy et al., 2012; Chen et al., 2016; Krashes et al., 2011), as well as generate foraging behavior in the absence of food (Dietrich et al., 2015; Krashes et al., 2011; Padilla et al., 2016). Previous work has begun to reveal the mechanisms by which motivational signals from the AgRP neurons can impact decision making. For example, a disynaptic circuit from AgRP neurons to posterior BNST is responsible for inhibiting territorial aggression during feeding (Padilla et al., 2016). It has also been shown that AgRP neurons are activated in response to social interactions and presentation of conspecifics when mice are hungry (Burnett et al., 2016). Our results suggest a novel mechanism that allows AgRP signaling to influence choice behavior via BNST^Y1R^ activation.

To survive, organisms must maintain essential physiological variables within an acceptable range. A signal representing energy deficit can be used to drive food seeking such that feeding is initiated when there is a deficit and halted when the energy supply has been replenished. AgRP neurons are often thought to provide a homeostatic error signal that is reduced by satiety and increased by deprivation. When reward is defined as error reduction in such a control system, then there must be temporal or delay discounting (Keramati and Gutkin, 2014). For example, in a system that controls for the rate of food intake, the error signal will be proportional to the degree of discounting. Larger errors will simply result in a higher rate of intake, leading to a preference for immediate rewards. Our results are in agreement with this possibility.

We showed that AgRP projections to the BNST are responsible for the observed effects on discounting. These projections release NPY and activate Y1Rs in BNST neurons. Indeed, BNST^Y1R^ activation, which is known to enhance inhibitory synaptic transmission (Pleil et al., 2015), is sufficient to restore delay discounting (**Figure 7**). AgRP neurons are known to co-release GABA (Tong et al., 2008), and GABA release from AgRP neurons is critical for feeding (Tong et al., 2008; Wu et al., 2009). Activation of NPY can also increase GABAergic signaling (Pleil et al., 2015), supporting the hypothesis that AgRP projections release NPY to reduce the activity of a subset of postsynaptic BNST-Y1R-expressing neurons. Our results are in agreement with this and demonstrate a novel pathway from AgRP neurons to BNST for the control of delay discounting in decision making.

The BNST has been implicated in stress, anxiety, and other aversive motivational states (Marcinkiewcz et al., 2016; Sahuque et al., 2006). Interestingly, AgRP activation suppress anxiety-like behavior (Burnett et al., 2016) and territorial aggression via projections to the BNST (Padilla et al., 2016). Our results thus link a key node in the homeostatic feeding system with circuitry known to control aversive motivational states. This link is also supported by previous work showing that activation of the stress-related BNST circuit results in weight loss (Kocho-Schellenberg et al., 2014). It is possible that, by suppressing aversive states, NPY signaling in the AgRP→BNST pathway can bias decision strategies and promote immediate gratification. Future work will be needed to further elucidate such appetitive-aversive interactions.

We have shown that the same neural circuit elements that are known to orchestrate feeding and regulate energy homeostasis are capable of biasing choice behavior and influencing self control. While AgRP neurons are rarely studied in animal models of psychiatric disorders, the present results demonstrate that they can influence high-level decision making processes via projections to the extended amygdala. Future studies may explore this pathway as a potential therapeutic target for disorders involving both food and decision making (e.g., anorexia nervosa or binge eating disorder).

### Experimental methods

#### Subjects

For behavioral experiments, Ai32 mice (Madisen et al., 2012) were crossed with AgRP-IRES-Cre mice (Tong et al., 2008), yielding mice with selective expression of ChR2 in AgRP-positive neurons (AgRP::ChR2; 3-7 months). Controls were ChR2-negative AgRP-Cre littermates. For electrophysiological experiments, AgRP::ChR2 mice (n = 5, 6-7 months, 2 male) were used. For drug infusion experiments, mice were all wild-type c57bl6 (n = 6, 3 months, all males). All experiments were approved by the Duke University Institutional Animal Care and Use Committee and were conducted in accordance with the National Institutes of Health guidelines regarding the care and use of animals.

#### Bilateral fiber optic implant surgery

Fiber optic implants were constructed as previously described (Rossi et al., 2012). Mice were anesthetized with isoflurane. For AgRP cell body stimulation group, craniotomies were made at −1.8 mm AP and +/−1.6 mm ML from Bregma. Fibers were then lowered 5.3 mm from the brain's surface at an angle of 14 ͦ relative to the vertical plane and fixed in place with dental acrylic and skull screws. For AgRP-BNST terminal stimulation group, craniotomies were made at −0.3 mm AP and +/−1.1 mm ML from Bregma. Fibers were then lowered 4.1 mm from the brain's surface at an angle of 6 ͦ. Mice were allowed to recover for at least one week before testing began.

#### Chronic cannulae implants surgery for drug infusions

Mice were anesthetized with isoflurane, and holes were drilled in the skull at −0.3 mm AP and +/−1.5 mm ML from Bregma. Cannulae (Plastic One, part number 8IC315GS4SPC) were then lowered 4 mm from the brain's surface at an angle of 10 ͦ and fixed in place with dental acrylic and skull screws. Finally, cannula dummies (Plastic One, part number 8IC315DC4SP) were fastened onto the cannulae to prevent clogging. Mice were allowed to recover for at least two weeks before testing began.

#### Temporal discounting behavior

Mice were group housed (2-5 per cage), and water was always available in the home cage. All testing took place in operant chambers (Med Associates) designed for in vivo optogenetics as described previously (Rossi et al., 2013). The chambers had two retractable levers located in one wall and a food cup located between the levers.

For the 1 pellet (same reward size both choices) temporal discounting task, the start of a trial was signaled by the extension of both levers. Mice could press either lever. Immediately following a lever press, both levers were retracted. For sessions in which a delay was imposed, one lever was designated the ‘delayed’ lever, and the other, the ‘immediate’ lever (lever designation was counterbalanced across mice). When mice pressed the immediate lever, a single food pellet (14 mg rodent purified diet, Bio Serv) was immediately delivered into the food cup. When mice pressed the delayed lever, a pellet was delivered into the food cup at a delay (4, 8, or 16). For the 0s delay condition, pressing either lever yielded immediate delivery of 1 food pellet. The following trial began 20 s after either lever was pressed. Only one delay condition was present during a session.

For the 4 pellet (larger reward for delay choice) task, pressing the immediate lever earned 1 pellet, but pressing the ‘delayed’ lever earned 4 pellets instead of 1 pellet. During the 0s delay condition, 1 lever was randomly designated the ‘delayed’ lever. The procedures are otherwise identical to the 1 pellet condition.

During initial lever press training, both levers yielded immediate delivery of 1 pellet. Following acquisition of lever pressing, temporal discounting sessions began. Sessions lasted 60 minutes and were self-paced. During testing, mice were hungry, sated, or pre-fed. When hungry, food was restricted such that they were maintained at ∼85-90% of free-feeding body weight. When sated, food was given *ad libitum* for 2-4 days prior to behavioral testing. When pre-fed (only in the first behavioral experiment), mice were allowed to consume 1.5 g of the same food pellets immediately prior to testing.

Mice were trained for 5-8 daily sessions on each delay. The delay was progressively increased (from 0 s to 16 s). To control for biases in lever preference, the delay and non-delay levers were switched for each delay condition. Stimulation or drug infusion experiments were performed once their performance had reached steady state on a given delay (4-7 days). The order of different stimulation parameters and drug concentrations was counterbalanced. For sessions in which mice failed to eat more than ∼20-30 of the pellets earned, the session was re-ran until mice consumed at least approximately 90% of pellets earned.

#### Optogenetic stimulation

All optogenetic stimulation was delivered bilaterally. Prior to testing, mice were connected to a 473 nm DPSS laser that delivered ∼10 mW (∼318 mW/mm^2^) into each hemisphere, as previously described (Rossi et al., 2012). For the entire session, laser pulse trains were delivered in 1 s bursts of square pulses (10 ms pulse width) at 10 Hz, 20 Hz, or 40 Hz followed by 3 s with no stimulation (Aponte et al., 2011). The order of stimulation was randomized. The lasers were controlled by a Blackrock data acquisition system.

#### Drug infusions

Drug infusions were administered bilaterally prior to behavioral testing. For [Leu^31^,Pro^34^]-Neuropeptide Y (*Tocris Cat. No. 1176*) infusions, pre-weighed drug (0.125 mg/kg, 0.25 mg/kg) dissolved in 500 nL phosphate-buffered saline (PBS) was infused into each hemisphere. For vehicle infusions, 500 nL PBS was infused into each hemisphere. Mice were first anesthetized with isoflurane, and then the drug or vehicle was delivered by a dual infusion pump (*Harvard apparatus. Cat. No* 13612) at 100 nL/min infusion rate. After infusions, mice were left in their home cage for 20 minutes for recovery before testing. The order of drug/vehicle infusions was randomized between mice.

#### In vitro electrophysiology

For whole cell patch clamp recordings, coronal sections were sliced from AgRP::ChR2 mice (n = 5). Brains were isolated quickly and immediately drowned into ice-cold solution bubbled with 95% O2-5% CO_2_ containing the following (in mM): 194 sucrose, 30 NaCl, 2.5 KCl, 1 MgCl2, 26 NaHCO3, 1.2 NaH2PO4, and 10 D-glucose with pH adjusted to 7.4 with HCl and osmolarity set to ∼320 mosM. After 5 minutes coronal slices were taken at 250 µm. During the recovery period (∼1 hr) slices were left in 35°C artificial cerebrospinal fluid (aCSF) solution bubbled with 95% O2-5% CO_2_ containing the following (in mM): 124 NaCl, 2.5 KCl, 2 CaCl2, 1 MgCl2, 26 NaHCO3, 1.2 NaH2PO4, and 10 D-glucose with pH adjusted to 7.4 with HCl and osmolarity set to ∼320 mosM. Following recovery, recorded under continuous perfusion of aCSF at 29-30°C.

For voltage-clamp experiments, the internal solution contained the following (in mM): 120 cesium methane sulfonate, 5 NaCl, 10 tetraethylammonium chloride, 10 HEPES, 4 lidocaine N-ethyl bromide, 1.1 EGTA, 4 magnesium ATP, and 0.3 sodium GTP, pH adjusted to 7.2 with CsOH and osmolarity set to 298 mosM with sucrose. For current-clamp experiments, the internal solution contained (in mM) 150 potassium gluconate, 2 MgCl2, 1.1 EGTA, 10 HEPES, 3 sodium ATP, and 0.2 sodium GTP, with pH adjusted to 7.2 with KOH and osmolarity set to ∼300 mosM with sucrose. Pipettes impedances were between 3.5 and 5 MΩ.

Slices were stimulated with 470-nm light generated from an LED (Thor Labs) focused through a ×40 objective (Olympus). During recordings, 10-ms flashes of light were delivered at 10 – 40 Hz to the entire ×40 field with an LED current driver (Thor Labs). Power density was ∼5 mW/mm^2^.

All data were recorded by a MultiClamp 700B amplifier (Molecular Device) and filtered at 10 kHz and digitized at 20 kHz with a Digidata 1440A digitizer (Molecular Devices). In the whole-cell configuration, recording was only accepted when the series resistance is < 20 MΩ. All data were analyzed using peak detection software in pCLAMP10 (Molecular Devices).

#### Anatomy and histology

Mice were perfused with 4% paraformaldehyde and brains post-fixed for 24 hours in 30% sucrose prior to vibratome sectioning at 60 mm coronally. Fiber and cannulae implantation sites were verified after sections were processed for the presence of cytochrome oxidase to visualize cytoarchitecture by rinsing in 0.1M PB before incubating in a diaminobenzidine, cytochrome C, and sucrose solution for ∼2 hours at room temperature. Mounted cytochrome oxidase sections were then dehydrated in 200 proof ethanol, defatted in xylene, and coverslipped with cytoseal. To confirm eYFP expression in AgRP cells in the ARC of AgRP-cre transgenic mice, select sections were rinsed in 0.1M PBS for 20 min before being placed in a PBS-based blocking solution containing 5% goat serum and 0.25% Triton X-100 at room temperature for 1 hr. Sections were then incubated with a primary antibody (polyclonal rabbit anti-AgRP; 1:200 dilution; ThermoFisher; catalog no. PA5-78739) in blocking solution overnight at 4 °C. Sections were then rinsed in PBS for 20 min before being placed in a secondary antibody used to visualize AgRP neurons in the ARC (goat anti-rabbit Alexa Fluor 594; 1:500 dilution; abcam; catalog no. ab150080) for 1 hr at room temperature. Visualization of NPY1R neurons in the BNST followed the same immunohistochemistry protocol with a different primary antibody (polyclonal rabbit anti-NPY1R; 1:200 dilution; Sigma-Aldrich; catalog no. SAB2101626). Sections for fluorescent microscopy were mounted and immediately coverslipped with Fluoromount G with DAPI medium (Electron Microscopy Sciences; catalog no. 17984-24). Brightfield images for placement were acquired and stitched using an Axio Imager M1 upright microscope (Zeiss) and fluorescent images were acquired and stitched using a Z10 inverted microscope (Zeiss).

#### Data analysis

Behavioral data were analyzed with Graphpad Prism. Choice on the delay lever was normalized to baseline lever preference with zero delay. Normalized choice preference was fit with a hyperbolic model (Doya, 2008):

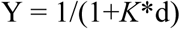

Where the value associated with the choice (Y) varies as a function of delay, d, and the discounting parameter, *K*, which determines the steepness of discounting.

For all behavioral data, the appropriate one-way or two-way ANOVA tests and post hoc comparisons were performed.

## Author Contributions

MAR and HHY conceived and designed the study. MAR, HEL, GWW, HGM, MTC, NK, KV, DL, and RAB conducted the experiments. MAR and HHY wrote the manuscript.

## Acknowledgments

This research was supported by a NSF fellowship to MAR. We would like to thank Tatyana Sukharnikova, Erin Gaidis, and Guozhong Yu for their help with histology.

## Supplementary materials

**Extended Figure 1.**
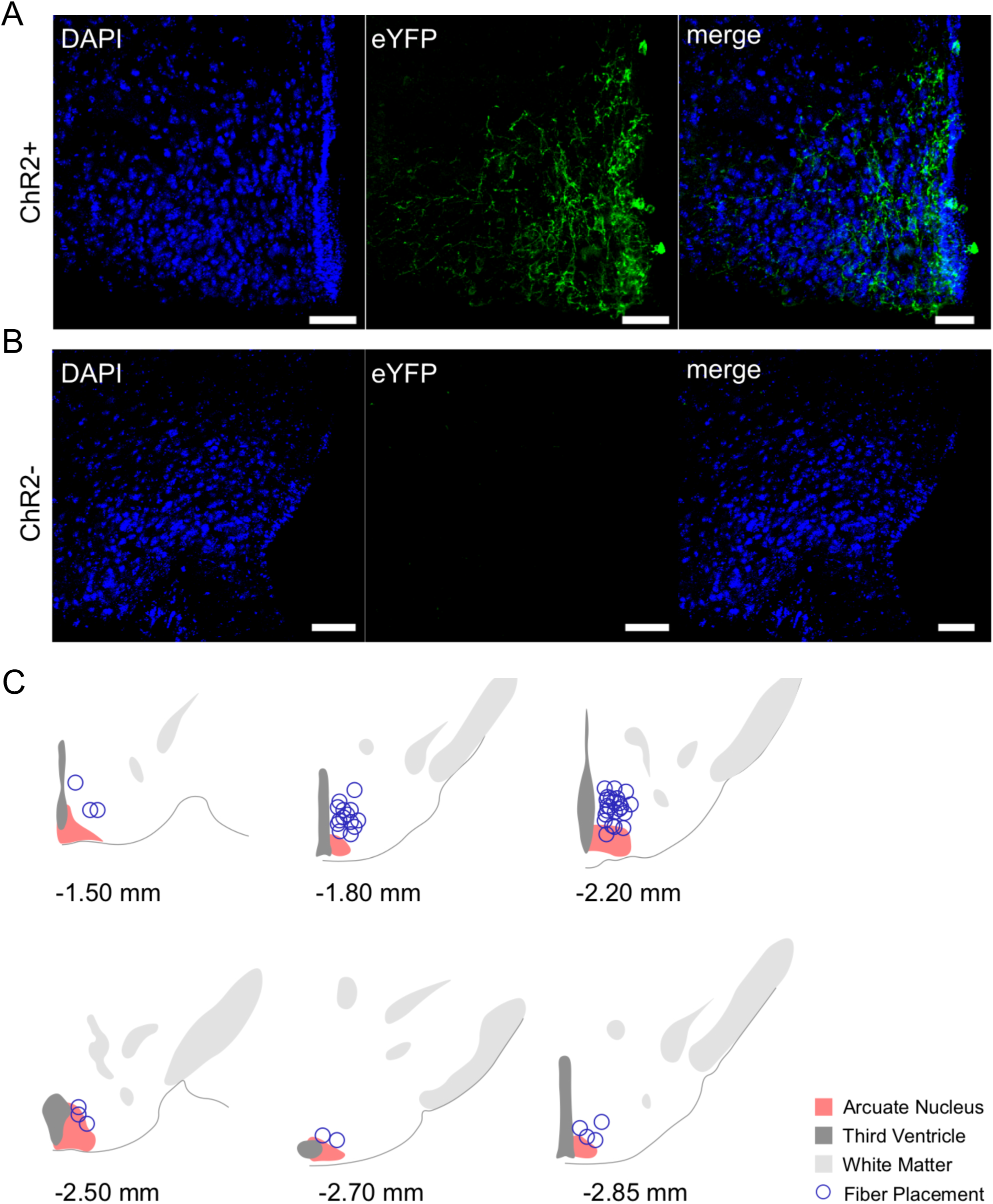
Opsin expression and optic fiber placements in the arcuate nucleus. **A.** Representative eYFP expression in AgRP::ChR2-eYFP mice. Scale bars, 50 µm. **B.** ChR2-littermates (AgRP-Cre control mice) have no detectable ChR2 expression. **C.** Placements of AgRP::ChR2-eYFP optical fibers and AgRP-Cre control optical fibers targeting ARC.

**Extended Figure 2.**
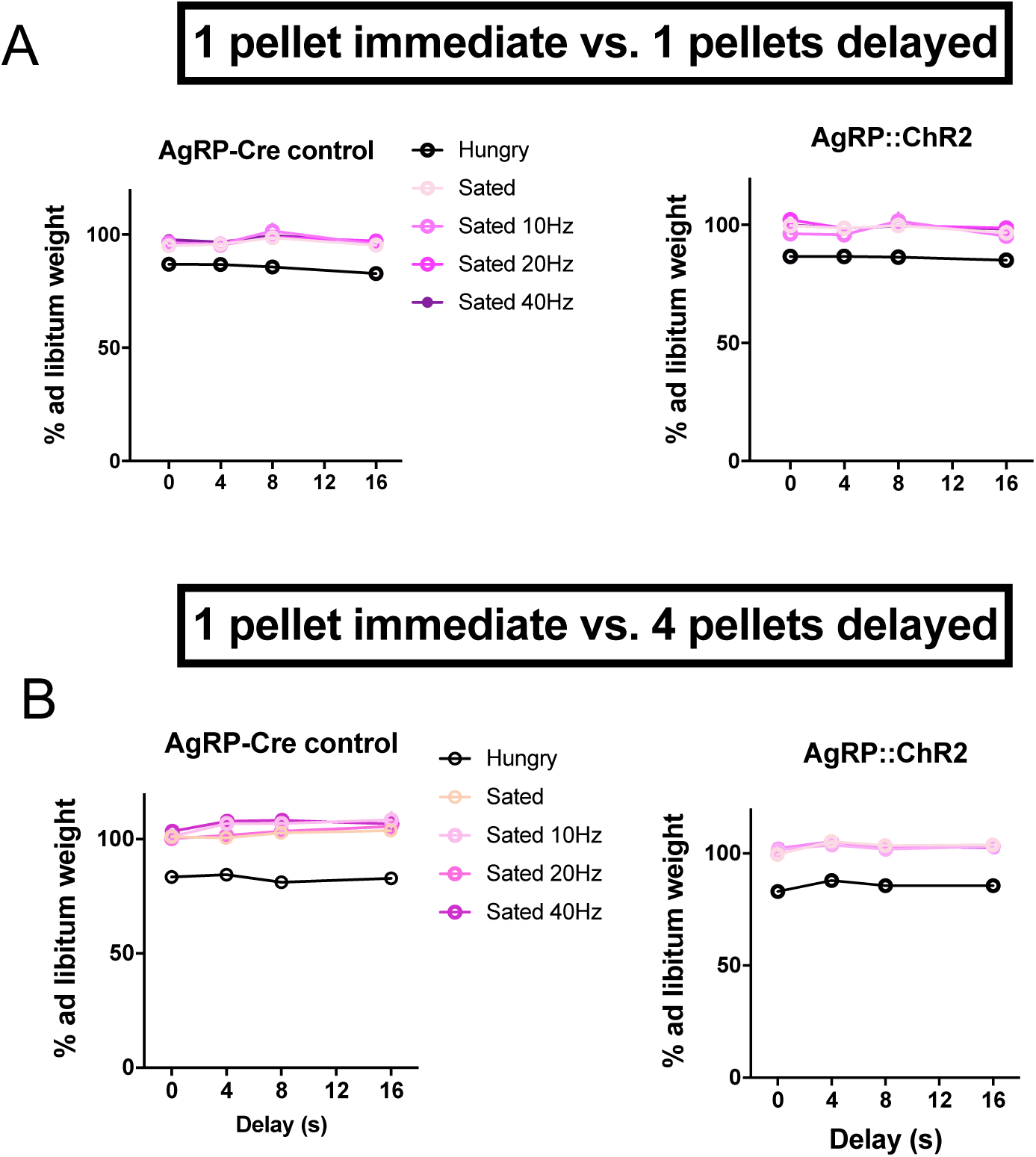
Body weights across stimulation conditions for AgRP cell body stimulation experiments. Weight was similar for all sated tests but reduced during Hungry tests. Error bars indicate ± s.e.m. **A.** Body weights for experiments in which the immediate and delayed rewards were the same magnitude. **B.** Body weights for experiments in which the delayed reward was larger than the immediate reward.

**Extended Figure 3.**
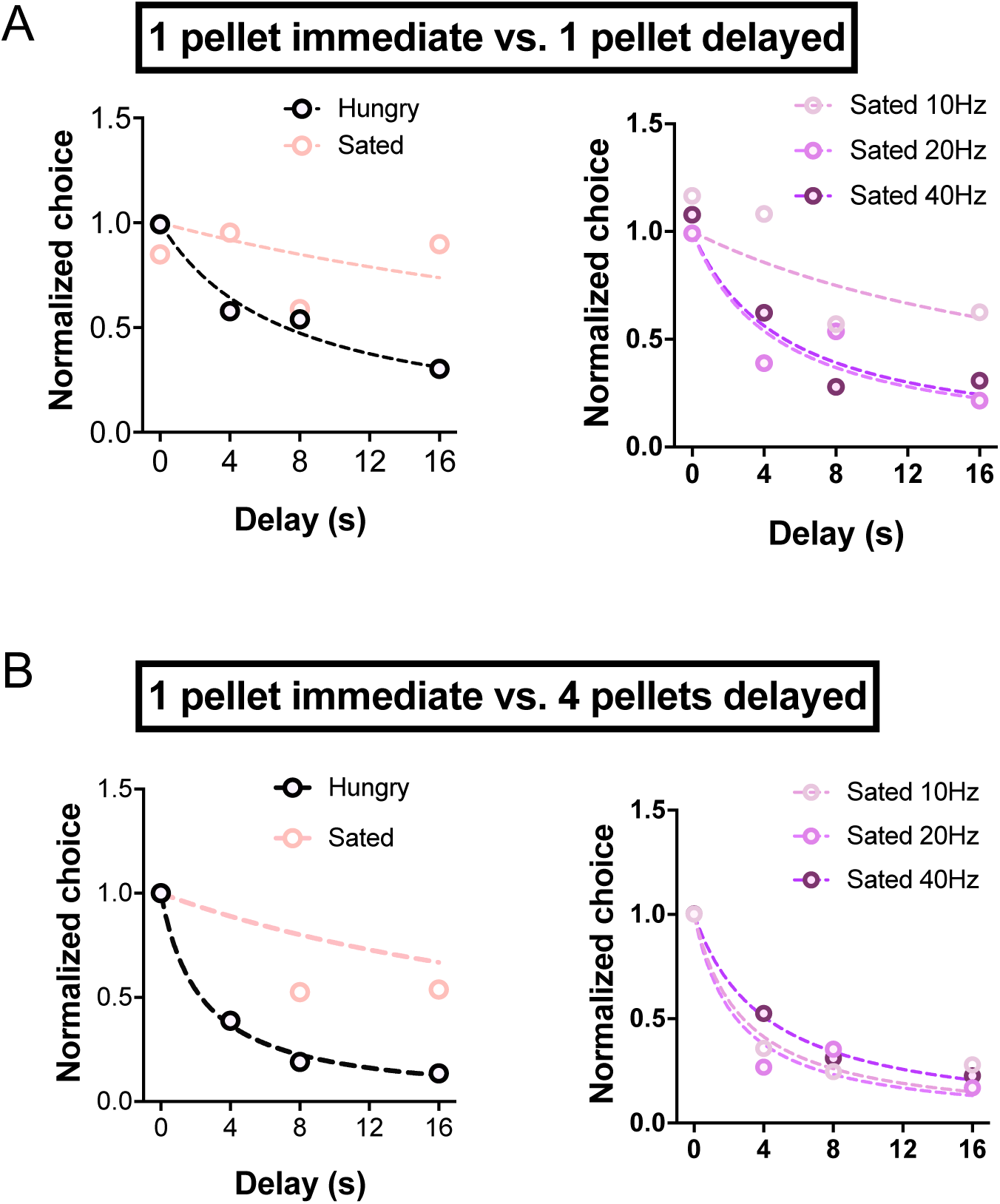
Representative examples of AgRP cell body stimulation. **A.** Temporal discounting functions of a representative AgRP::ChR2-eYFP mouse in the pure delay discounting condition, in which the delayed reward is the same size as the immediate reward (1 pellet). Dashed lines represent hyperbolic fits. **B.** Discounting functions for the delay/larger reward (4 pellets) condition.

**Extended Figure 4.**
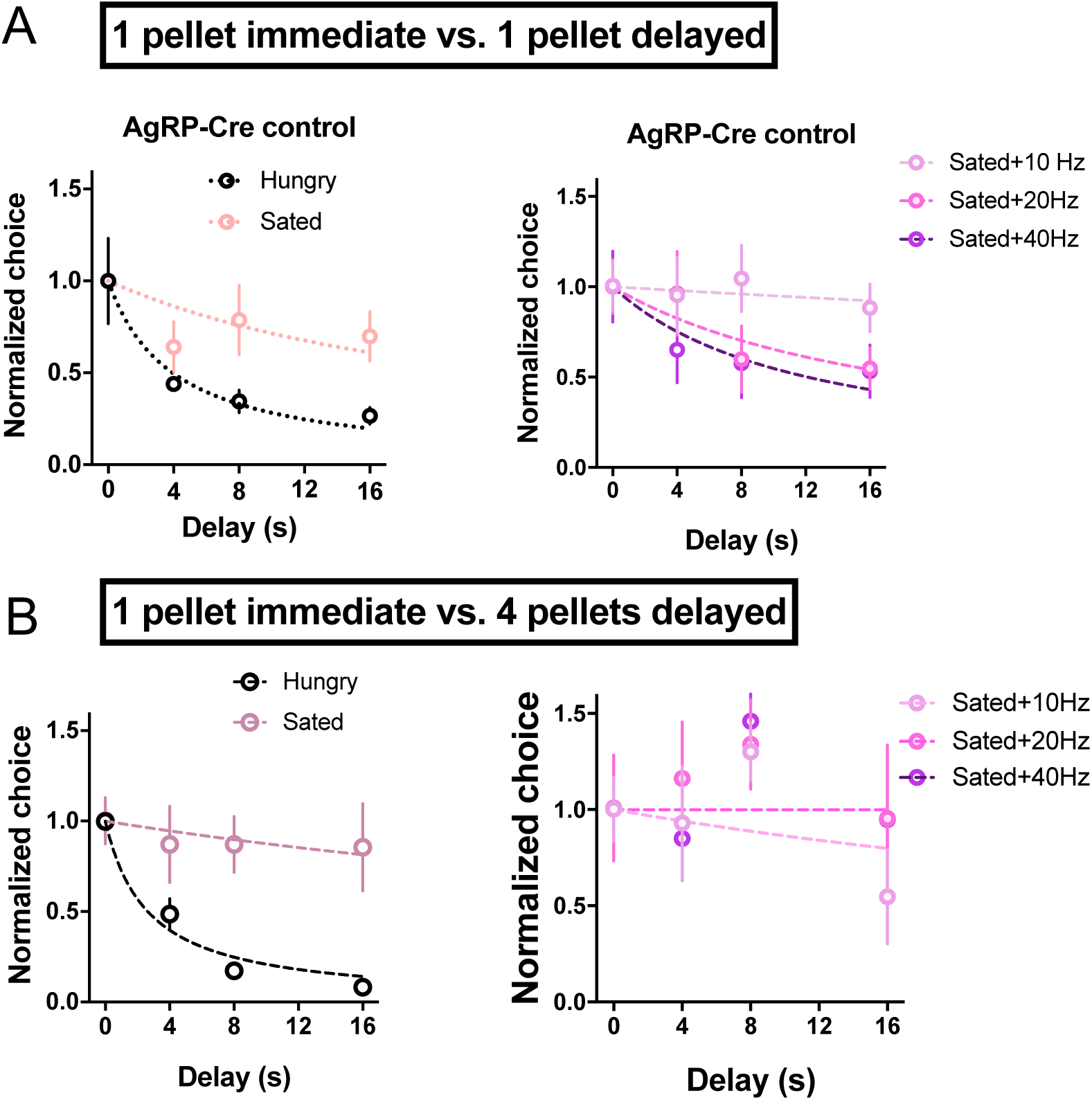
Discounting is not affected by AgRP cell body stimulation in control mice. **A.** Discounting functions for AgRP-Cre control mice in the pure delay discounting condition, in which the delayed reward is the same size as the immediate reward (1 pellet). Dashed lines represent hyperbolic fits. **B.** Discounting functions for the delay/larger reward (4 pellets) condition.

**Extended Figure 5.**
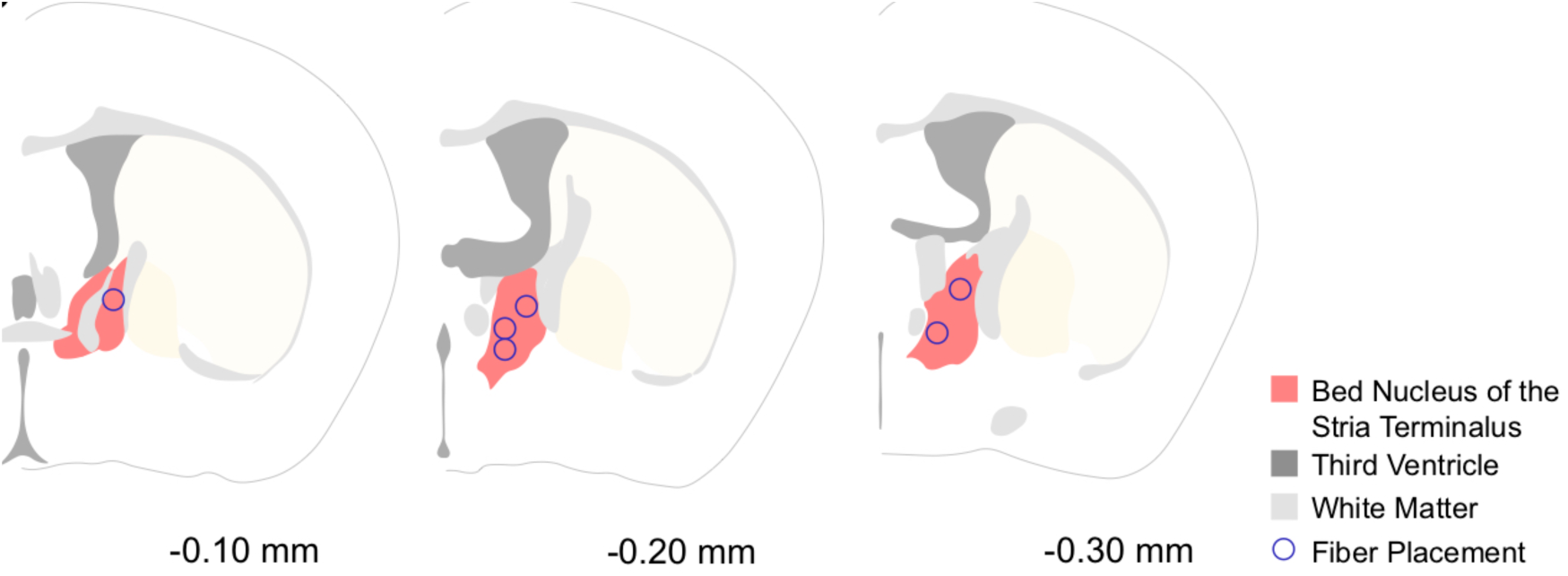
Optic fiber placements in the BNST from AgRP::ChR2→BNST terminal stimulation experiments.

**Extended Figure 6.**
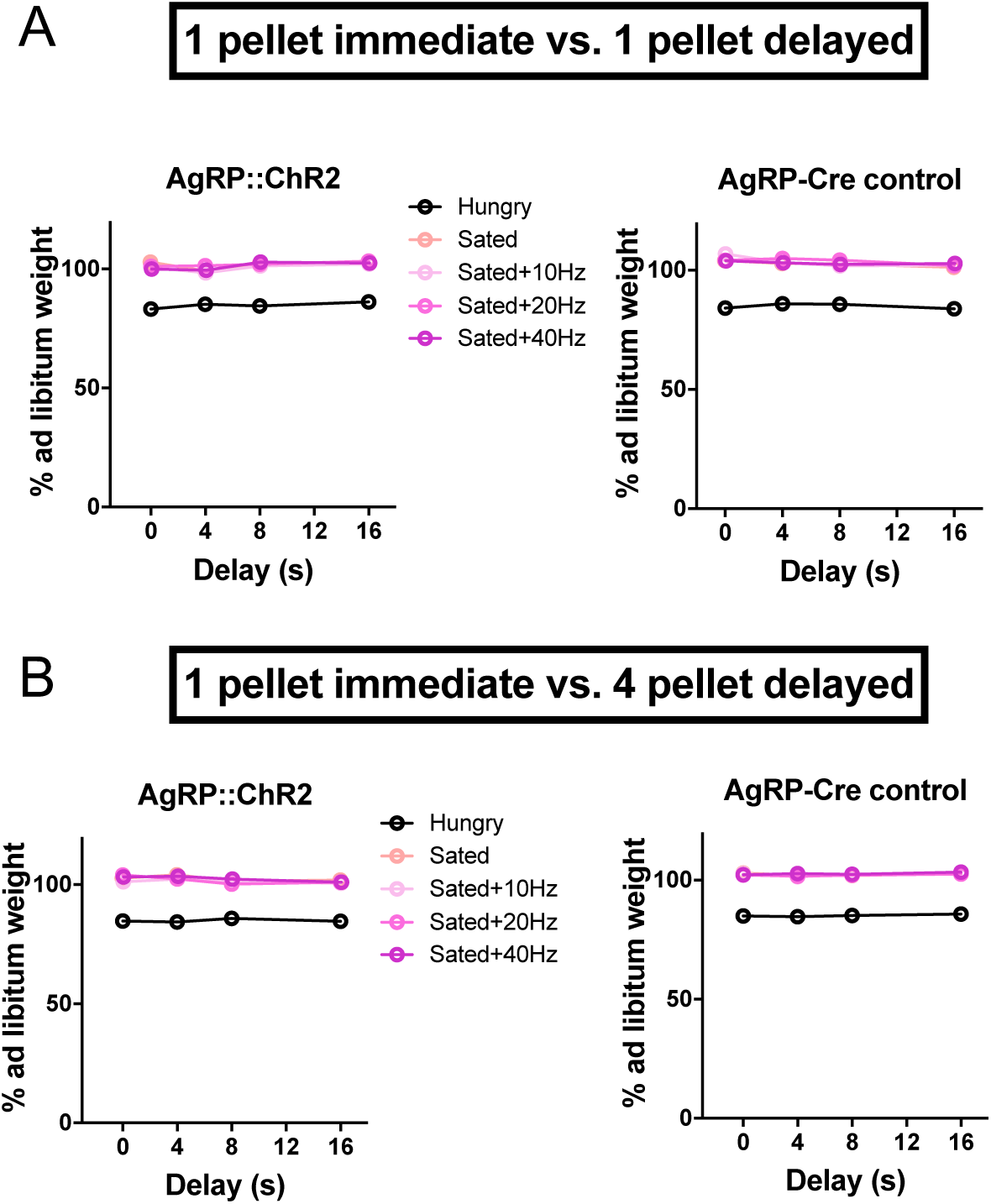
Body weights across stimulation conditions for AgRP::ChR2→BNST terminal stimulation experiments. Weight was similar for all sated tests but reduced during Hungry tests. Error bars indicate ± s.e.m. **A.** Body weights for experiments in which the immediate and delayed rewards were the same magnitude. **B.** Body weights for experiments in which the delayed reward was larger than the immediate reward.

**Extended Figure 7.**
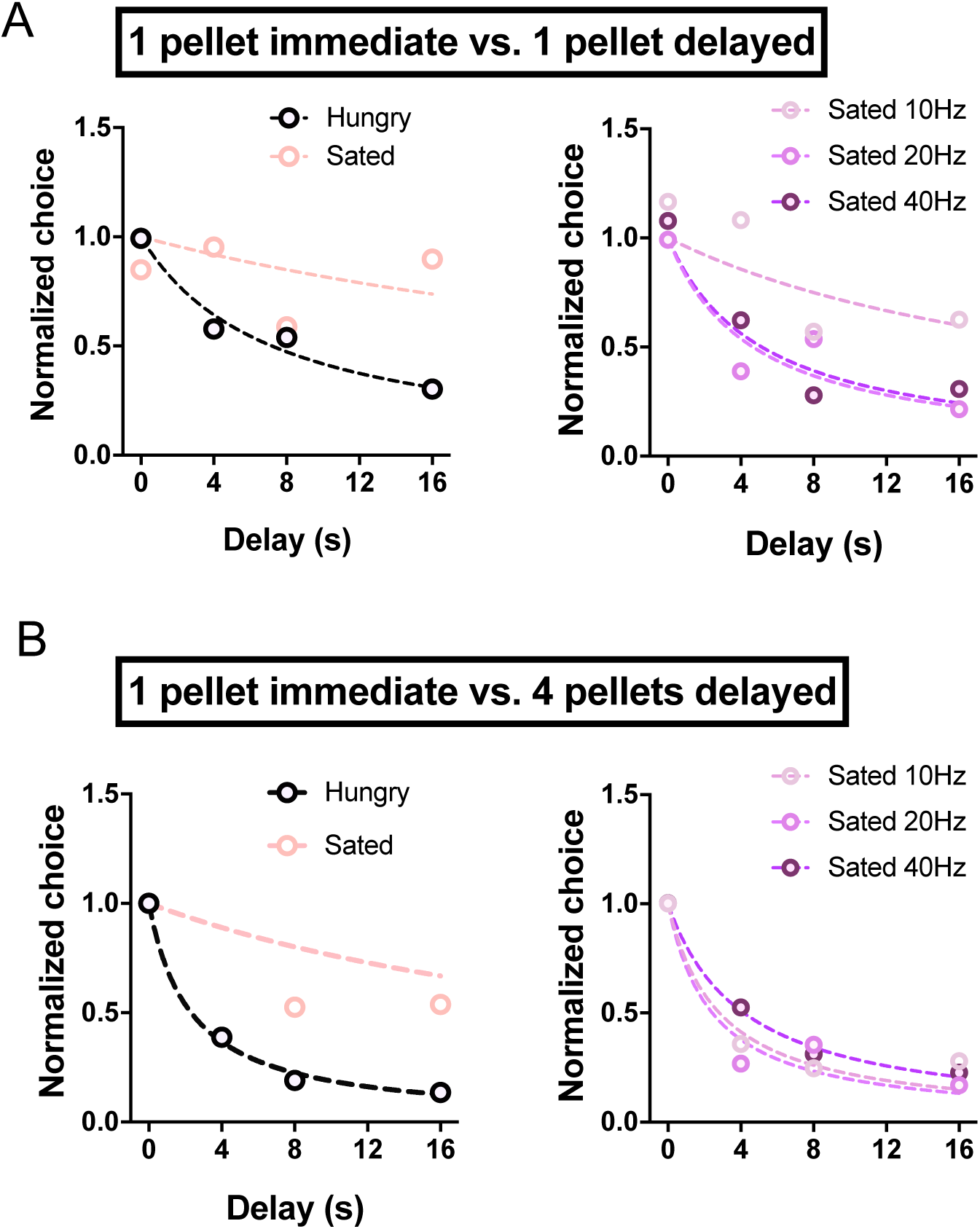
Representative examples of AgRP::ChR2→BNST terminal stimulation. **A.** Discounting function of a representative mouse with AgRP::ChR2→ stimulation in the pure delay discounting condition, in which the delayed reward is the same size as the immediate reward (1 pellet). Dashed lines represent hyperbolic fits. **B.** Discounting function of a representative mouse for the delay/larger reward (4 pellets) condition.

**Extended Figure 8.**
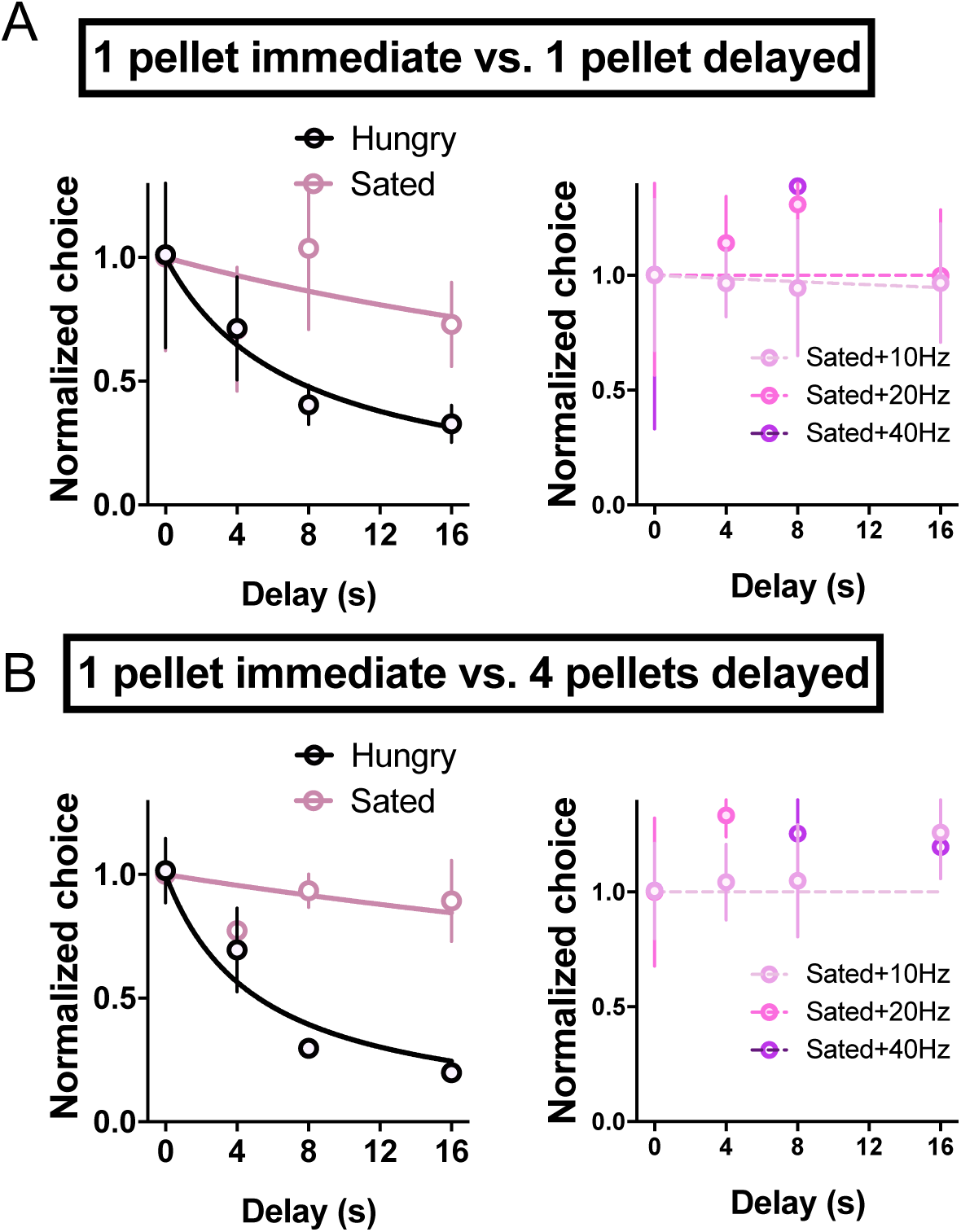
Discounting is unaffected by AgRP::ChR2→BNST terminal stimulation in control mice. **A.** Discounting functions for AgRP-Cre control mice in the pure delay discounting condition, in which the delayed reward is the same size as the immediate reward (1 pellet). Dashed lines represent hyperbolic fits. **B.** Discounting functions for the delay/larger reward (4 pellets) condition.

**Extended Figure 9.**
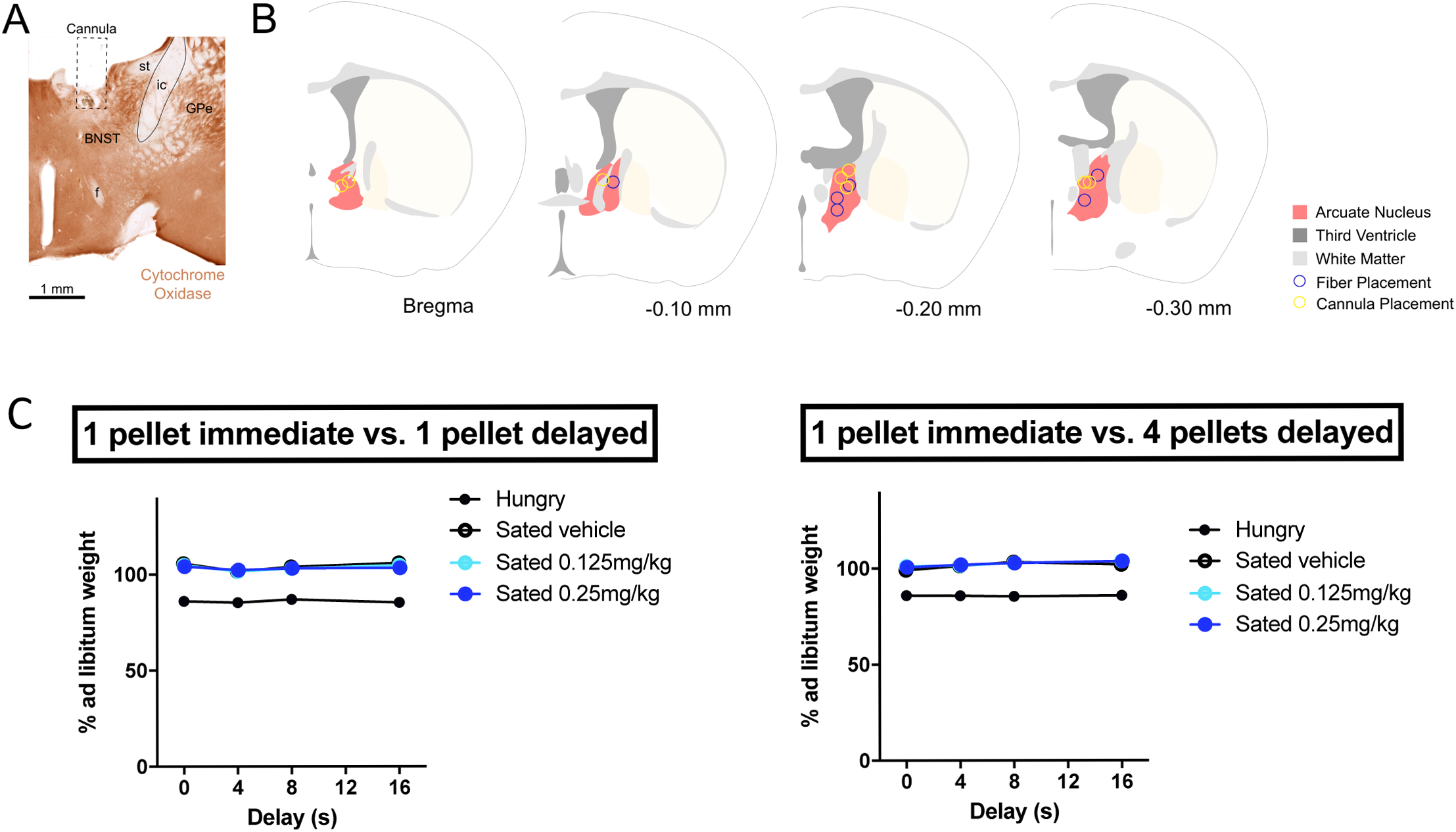
Cannula placement in the BNST for Leu-Pro NPY injection experiments. **A.** Cytochrome oxidase staining (coronal section) showing representative cannula placement **B.** Illustration of cannula placement in all mice tested. **C.** Body weights of mice from the Leu-Pro NPY injection experiments. Error bars indicate mean ± s.e.m.

**Extended Figure 10.**
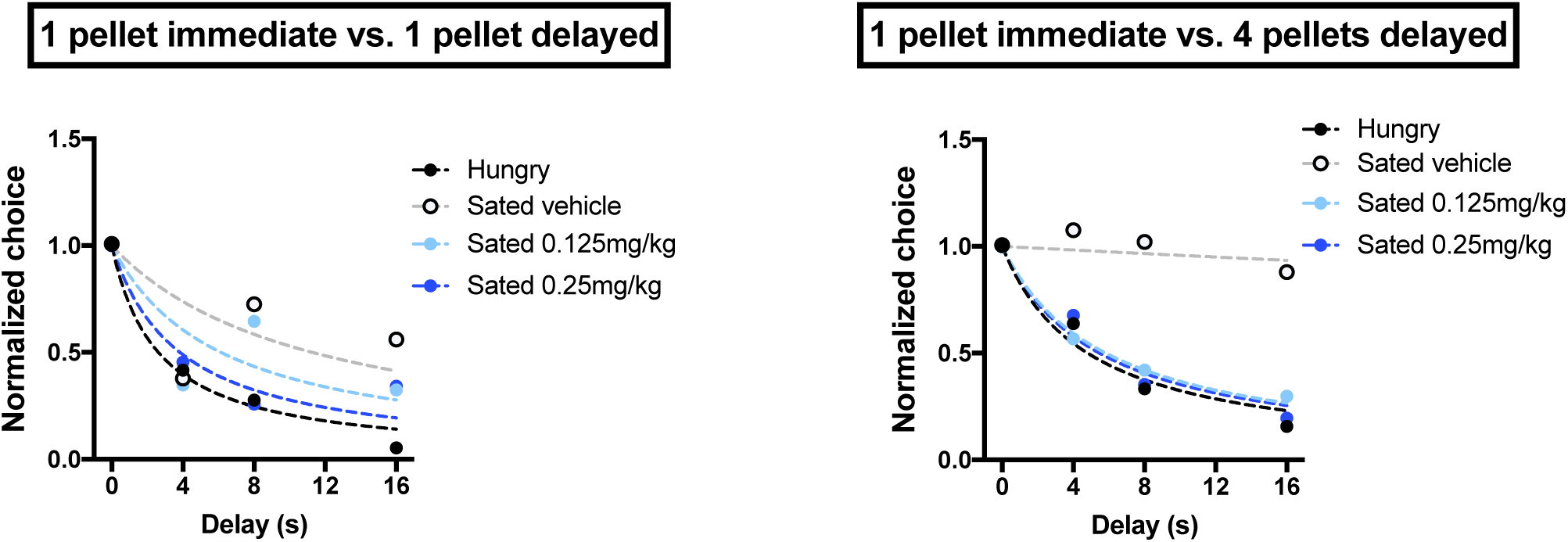
Representative temporal discounting functions from Leu-Pro NPY injection experiments.

